# ATR safeguards replication forks against APOBEC3B-induced toxic PARP1 trapping

**DOI:** 10.1101/2024.11.14.623607

**Authors:** Pedro Ortega, Elodie Bournique, Junyi Li, Ambrocio Sanchez, Gisselle Santiago, Brooke R. Harris, Abby M. Green, Rémi Buisson

## Abstract

ATR is the master safeguard of genomic integrity during DNA replication. Acute inhibition of ATR with ATR inhibitor (ATRi) triggers a surge in origin firing, leading to increased levels of single-stranded DNA (ssDNA) that rapidly deplete all available RPA. This leaves ssDNA unprotected and susceptible to breakage, a phenomenon known as replication catastrophe. However, the mechanism by which unprotected ssDNA breaks remains unclear. Here, we reveal that APOBEC3B is the key enzyme targeting unprotected ssDNA at replication forks, triggering a reaction cascade that induces fork collapse and PARP1 hyperactivation. Mechanistically, we demonstrate that uracils generated by APOBEC3B at replication forks are removed by UNG2, creating abasic sites that are subsequently cleaved by APE1 endonuclease. Moreover, we demonstrate that APE1-mediated DNA cleavage is the critical enzymatic step for PARP1 trapping and hyperactivation in cells, regardless of how abasic sites are generated on DNA. Finally, we show that APOBEC3B-induced toxic PARP1 trapping in response to ATRi drives cell sensitivity to ATR inhibition, creating to a context of synthetic lethality when combined with PARP inhibitors. Together, these findings reveal the mechanisms that cause replication forks to break during replication catastrophe and explain why ATRi-treated cells are particularly sensitive to PARP inhibitors.

## INTRODUCTION

The ataxia telangiectasia and Rad3-related (ATR) checkpoint kinase is a critical safeguard of the genome. ATR directly controls DNA replication by maintaining proper levels of origin firing during Sphase ^1–3^. Moreover, ATR is recruited to damaged replication forks, where it orchestrates replication fork repair and restart ^1,3^. The essential role of ATR in maintaining genome integrity makes it an attractive target for cancer therapy. With many ATR inhibitors (ATRi) currently in clinical trials ^4^, it is crucial to understand the cellular contexts that render cancers sensitive to ATRi. Cancer cells with defects in DNA repair pathways, such as homologous recombination, or experiencing replication stress driven by specific oncogenes, become particularly vulnerable to ATRi ^5–8^. Notably, cells treated with ATRi become hypersensitive to PARP inhibitors (PARPi) ^7,9^, an enzyme that detects DNA damage and facilitates the recruitment of DNA repair proteins through its poly ADP-ribosylation activity ^10,11^. However, it still remains unclear why PARP inhibition is so effective in killing cells without ATR activity.

ATR inhibition triggers a specific type of replication stress in cells through the upregulation of CDK1/2 and CDC7 activity ^12–14^, increasing origin firings and suppressing RRM2, a cell-cycle-regulated subunit of the ribonucleotide reductase critical for dNTP synthesis in the S-phase ^5,14–17^. Therefore, the increase in replication and decrease in dNTP synthesis upon acute ATR inhibition leads to the exhaustion of dNTPs, causing replication forks to slow down, uncoupling of helicases and polymerases, and accumulation of ssDNA associated with lagging strands of the replication forks^1–3^. Replication Protein A (RPA) is critical for protecting ssDNA at replication forks from nucleolytic attacks^18,19^. Following ATR inhibition, the surge of ssDNA levels depletes the pool of available RPA in cells, leaving ssDNA unprotected and susceptible to breakage, a phenomenon known as replication catastrophe ^5,19,20^. However, the mechanism by which unprotected ssDNA breaks at replication forks following ATR inhibition and RPA exhaustion is still poorly understood.

ATR is critical to safeguard cells from replication stress mediated by APOBEC3A (A3A) and APOBEC3B (A3B) deaminase activity, two members of the Apolipoprotein B mRNA-editing enzyme catalytic polypeptide-like (APOBEC) cytidine deaminase family. APOBEC enzymes are antiviral factors that promote the deamination of cytosine to uracil in DNA or RNA ^21,22^. APOBEC enzymes counteract the replication of various DNA or RNA viruses, retroviruses, and retrotransposons ^21,22^. In addition to their role in protecting cells against viral infection, APOBEC enzymes are one of the most common causes of genomic mutations in cancer ^23–28^. The APOBEC mutational signature is characterized by two distinct single-base substitutions (SBSs): SBS2 characterized by C>T mutations and SBS13 consisting of C>G and C>A substitutions, both occurring in TCA and TCT trinucleotide sequence contexts ^24,25,27^. APOBEC-signature mutations are preferentially enriched on the lagging-strand template of DNA replication forks ^29–31^. The strong association between APOBEC3 mutagenesis and the lagging strand of the replication forks underscores that transiently exposed single-stranded DNA (ss-DNA) during replication is the main target of APOBEC3 enzymes in cells.

A3A and A3B are responsible for most of the APOBEC mutational signatures identified in tumors ^32–38^. A3A expression in cancer cells is low and tightly regulated through the interferon response and the NF-kB pathway triggered by diverse cellular stresses encountered by cells, leading to episodic bursts of mutations ^39–41^. In contrast, A3B is highly expressed in many tumor types, including breast, lung, colorectal, bladder, cervical, head and neck, and ovarian cancers ^37,42,43^. A3B’s deaminase activity is less efficient than A3A, enabling cancer cells to tolerate constitutive and high expression levels ^37,44^. Aside from inducing mutations in cancer genomes, A3A and A3B also contribute to genomic instability by inducing replication stress, the formation of DNA double-strand breaks (DSBs), chromosomal instability (CIN), and aneuploidy ^36,45–49^. Given their ability to rewrite genomic information and increase genomic instability, A3A and A3B drive tumor diversity, promote cancer progression, and contribute to therapy resistance, all of which are associated with poor overall survival of the patients ^50–54^. Yet, it remains unclear how deaminated cytosines generated by APOBEC enzymes at replication forks lead to the formation of DSBs and genomic instability.

Cells expressing A3A or A3B are particularly sensitive to ATR inhibitors (ATRi) ^46,48,55^, suggesting a unique function of ATR in shielding replication forks against their deaminase activity. However, the mechanism by which ATR protects cells from A3A and A3B activity is still unclear. Previous studies, including from our laboratory, employed model cell lines ectopically overexpressing either A3A or A3B to study cells’ response to DNA damage caused by these enzymes ^38,46–48,52,54–59^. While these models have been essential to start understanding the consequences of APOBEC deaminase activity in cells, they raise concerns about how expressing an enzyme ten times or more than endogenous levels recapitulates their physiological roles. Indeed, high expression levels of A3A and A3B can potentially bypass normal cellular mechanisms that protect the genome from their DNA deaminase activity. Moreover, overexpression of A3A or A3B may activate different types of DNA repair processes that become necessary to respond to aberrant levels of deaminated cytosines but that are not normally required by cells damaged by endogenous levels of APOBEC3 enzymes. Therefore, it is critical to better understand the cellular response to APOBEC3 endogenous activity to effectively exploit in the future APOBEC3-induced cellular vulnerabilities to ATR inhibitors.

In this study, we identify A3B as the key enzyme promoting ssDNA breakage during replication catastrophe upon acute inhibition of ATR, leading to PARP1 hyperactivation. Mechanistically, we show that A3B targets unprotected ssDNA following RPA exhaustion caused by the surge of dormant origin firing mediated by ATR inhibition. Uracils generated by A3B at replication forks are then removed by UNG2 to form abasic sites that are subsequently cleaved by APE1, leading to replication fork collapse and PARP1 hyperactivation. Importantly, we explain why ATR inhibition is particularly effective at activating PARP1 compared to other types of DNA damage that cause fork collapse and DSB formation. Indeed, we reveal that APE1-mediated abasic cleavage is the key step driving PARP1 hyperactivation not only upon ATR inhibition but also following treatments that induce abasic sites, demonstrating that APE1 generates the specific substrate recognized by PARP1, regardless of how the abasic sites are formed. Finally, our results provide a mechanistic explanation for how A3B confers a therapeutic vulnerability to ATR inhibitors in combination with PARP1 inhibitors by mediating toxic PARP1 trapping at replication forks. Together, our findings suggest that A3B expression levels in cancer cells may serve as a potential biomarker for predicting the response to ATRi and PARPi therapies.

## RESULTS

### ATR prevents PARP1 hyperactivation during DNA replication

To investigate the unique cellular response to ATR inhibition that renders replication forks particularly susceptible to APOBEC deaminase activity, we first treated cells with a specific ATR inhibitor (ATRi; BAY-1895344) and compared the DNA damage signaling responses to other types of genotoxic stress. We selected hydroxyurea (HU) to block ongoing replication forks, the topoisomerase I inhibitor camptothecin (CPT) to generate ssDNA breaks leading to replication collapse, and the topoisomerase II inhibitor etoposide (ETP) to cause DSBs independently of DNA replication. All four treatments lead to ssDNA formation in damaged cells, which could be targeted by APOBEC enzymes. We next monitored different DNA damage markers, including H2AX, RPA, and Chk1 phosphorylation, as well as PARP MARylation and PARylation (MAR/PARylation) levels. Remarkably, we found that cells treated with ATRi elicited strong MAR/PARylation levels in cells compared to other DNA damage treatments **(Figure 1A and Supplementary Figure 1A)**, which was suppressed in cells that were knocked down for PARP1 or were treated with PARP inhibitor (PARPi) **(Figure 1B and Supplementary Figure 1B)**. We further confirmed this result using a different ATR inhibitor (ATRi#2; AZD6738) or by inhibiting the DNA damage checkpoint kinase Chk1 (Chk1i; MK-8776), the downstream target of ATR **(Figure 1C-D)**. Although DNA damages caused by HU, CPT, or ETP are known to activate PARP1^60–63^, the levels of MAR/PARylation following these treatments were very low compared to ATRi and were detectable only after treatment with PARG inhibitor (PARGi) to block PARG activity removing PAR chains on PARP1 or other proteins **(Figure 1E)**. Thus, these results suggest that ATR inhibition leads to a unique cellular response causing PARP1 hyperactivation.

**Figure 1:**
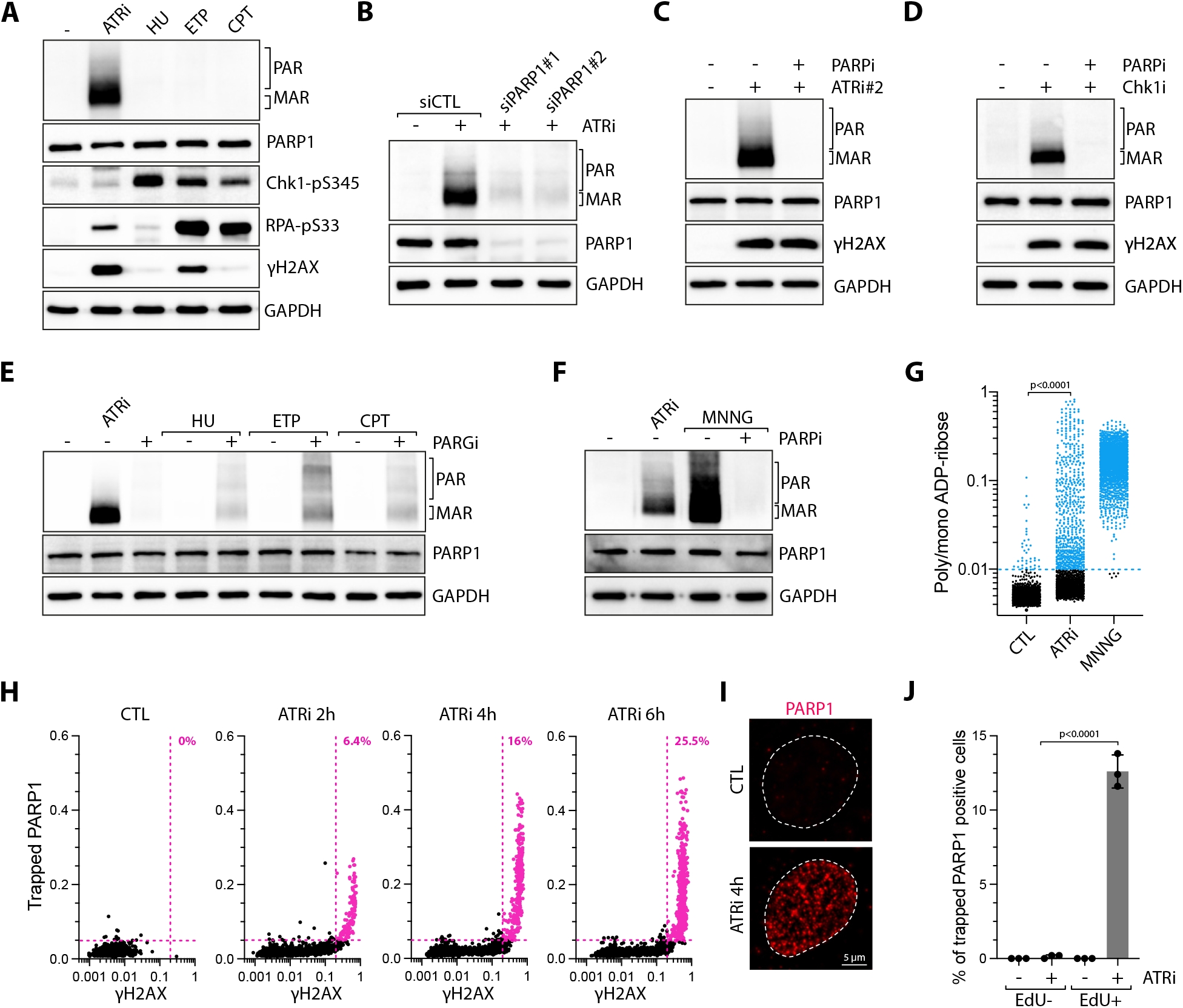
ATR inhibition induces PARP1 hyperactivation in S-phase. **A**. U2OS cells were treated with ATRi (1 μM), HU (2 mM), ETP (25 μM), or CPT (1 μM) for 4h and analyzed by western blot using the indicated antibodies. **B**. U2OS cells were transfected with siRNA against PARP1 for 40h and subsequently treated with ATRi (1 μM) for 4h, and the levels of MAR/PAR were analyzed by western blot. **C-D**. The levels of MAR/PAR were monitored by western blotting in U2OS cells treated with ATRi#2 (2 μM) **(C)** or Chk1i (1 μM) **(D)** for 4h in the presence or absence of PARPi (10 μM). **E**. U2OS cells were treated with ATRi (1 μM), HU (2 mM), ETP (25 μM), or CPT (1 μM) for 4h in the presence or absence of PARGi (2 μM). The levels of poly/mono ADP-ribose, PARP1, and GAPDH were analyzed by western blot. **F**. U2OS cells were treated with ATRi (1 μM; 4h) or MNNG (10 μM; 15 min) in the presence or absence of PARPi (10 μM with 1h pre-treatment), and the level of MAR/PAR was analyzed by western blot. **G**. Poly/mono ADPribose levels in the nucleus were monitored by immunofluorescence in 2,000 U2OS cells treated with ATRi (1 μM; 4h) or MNNG (10 μM; 1h). **H**. Quantification of nuclear PARP1 and γH2AX in 2,000 U2OS cells treated with ATRi (1 μM) for the indicated time. Colored dots and percentages indicate cells positive for γH2AX and PARP1. **I**. Representative immunofluorescence pictures for PARP1 staining in U2OS cells treated with DMSO or ATRi for 4h. Scale bar: 5 μm. **J**. Quantification of trapped PARP1-positive cells (%) across the whole population and subsequently categorized into EdU-positive or EdU-negative cells. U2OS cells were treated with EdU (10 μM) for 15 minutes before adding ATRi for 4 hours. Mean values ± SD (Number of biological replicates, n = 3). All P-values were calculated with a two-tailed Student t-test.

We then compared ATRi to MNNG (N-Methyl-N’nitro-N-nitrosoguanidine), a DNA alkylating agent well-characterized to trigger strong activation of PARP1 in cells ^64,65^. Cell treatment with MNNG strongly induces PARP1 MAR/PARylation **(Figure 1F)**. However, MNNG activates PARP1 in the whole cell population, while ATRi-induced PARP1 activation was confined to a fraction of cells **(Figure 1G)**. Nevertheless, cells positive for MAR/PARylation after ATRi displayed levels comparable to those in cells treated with MNNG **(Figure 1G)**, further demonstrating that ATR inhibition strongly predisposes a specific subset of cells to PARP1 hyperactivation. To better investigate PARP1 hyperactivation in response to ATR inhibition, we next employed a single-cell-based method using high-content microscopy to measure the levels of DNA damage and replication stress (γH2AX positive cells) as well as the amount of PARP1 associated with chromatin by pre-extracting cells with detergent before fixation to remove all soluble PARP1 **(Figure 1H)**. We found that ATRi treatment promotes a gradual increase in PARP1 trapping on chromatin **(Figure 1H)**. Moreover, we showed that PARP1 trapping and MAR/PARylation levels were associated with S-phase cells that were positive for γH2AX and EdU **(Figure 1H-J, and Supplementary Figure 1C)**. In contrast, MNNG treatment results in PARP1 trapping regardless of the cell cycle stages **(Supplementary Figure 1D)**. Taken together, these results reveal that ATR activity is critical to prevent hyperactivation of PARP1 in replicating cells.

### PARP1 hyperactivation recruits XRCC1 to replication forks

PARP1 is a DNA sensor enzyme that detects DNA breaks in cells and coordinates various cellular processes such as DNA repair, chromatin remodeling, and cell death ^10,11^. To investigate the role of PARP1 hyperactivation upon ATR inhibition in replicating cells, we first monitored PARP1 localization alongside the replication stress markers RPA and phosphorylated RPA, which are known to be recruited and activated at replication forks upon ATR inhibition ^5,20^. Following ATR inhibition, PARP1 formed distinct nuclear foci that colocalized with RPA and RPA-pS4/8 foci **(Figure 2A)**, further indicating PARP1’s association with damaged replication forks. Once recruited to DNA breaks, PARP1 initiates the MAR/PARylation on itself and other proteins to recruit DNA repair factors, stabilize the DNA damage site, and promote the repair process ^10,11^. Additionally, PARP1 can directly modify chromatin organization around the DNA breaks by facilitating the recruitment and regulation of chromatin remodelers ^66–70^. We next asked whether PARP1 activation at replication forks was required for the recruitment of specific DNA repair factors. Among the repair proteins localized at replication forks through the PARylation of PARP1 is XRCC1, a molecular scaffold important for the recruitment of many other repair proteins ^71,72^. We selected XRCC1 for our study not only due to its significance in repair processes at replication forks ^72–74^, but also because a specific antibody was available for its detection in cells via immunofluorescence ^75^. Similar to PARP1, XRCC1 was detected in γH2AX-positive cells following detergent extraction **(Figure 2B)**, indicating XRCC1’s association with damaged replicating cells. Importantly, XRCC1 formed discrete foci that colocalized with both RPA and PARP1 following ATRi treatment **(Figure 2C)**, indicating that XRCC1 is in complex with PARP1 at damaged replication forks. To test whether XRCC1 localization to damaged replication forks is mediated by PARP1, we monitored both XRCC1 and PARP1 association with chromatin upon ATRi or ATRi+PARPi treatments. While PARP1’s association with chromatin was not affected by PARPi, XRCC1 recruitment strongly decreased **(Figure 2D)**, demonstrating that PARP1 mediates XRCC1 recruitment to replication forks through its MAR/PARylation activity. Collectively, these results suggest that PARP1 hyperactivation in the absence of ATR activity is critical for the recruitment of PARP1-associated repair factors to damaged replication forks.

**Figure 2:**
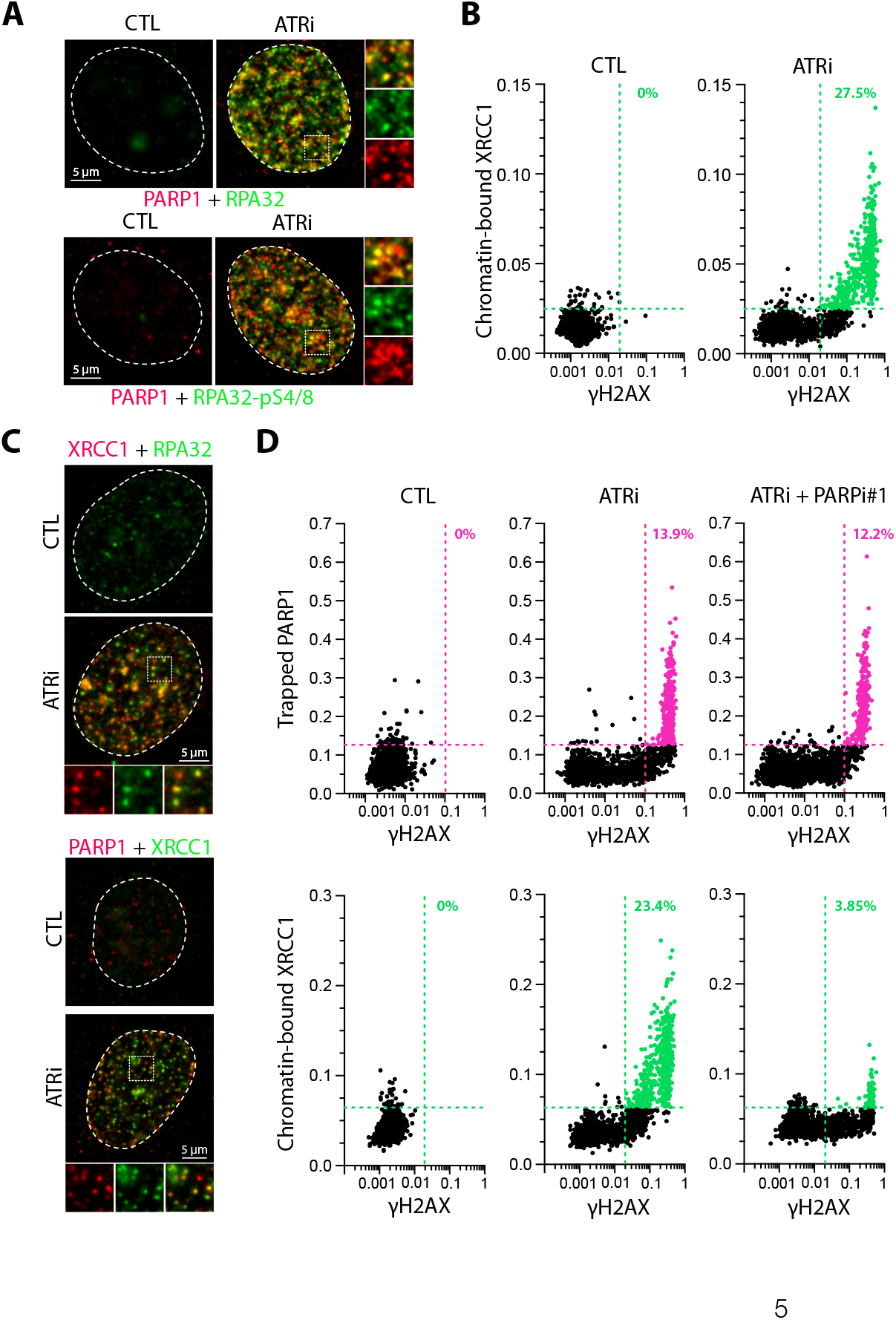
Recruitment of PARP1 and XRCC1 to replication forks upon ATR inhibition. **A**. U2OS cells were treated with ATRi (1 μM), and the cellular localization of PARP1, RPA32, or RPA32-pS4/8 was monitored by immunofluorescence. Scale bar: 5 μm. **B**. Quantification of chromatin-bound XRCC1 and γH2AX in 2,000 U2OS cells treated with ATRi (1 μM) for 4h. Colored dots and percentages indicate cells positive for γH2AX and PARP1. **C**. U2OS cells were treated with ATRi (1 μM), and the cellular localization of the indicated proteins was monitored by immunofluorescence. Scale bar: 5 μm. **D**. Quantification of nuclear PARP1 or XRCC1 and γH2AX in 2,000 U2OS cells treated with ATRi (1 μM) in the presence or absence of PARPi (20 μM) for 4h. Colored dots and percentages indicate cells positive for γH2AX and PARP1 or XRCC1.

### ATR suppresses PARP1 hyperactivation by limiting origin firings

To understand how ATR inhibition causes PARP1 hyperactivation at replication forks, we investigated which specific events mediated by ATR inhibitors are responsible for triggering the hyperactivation of PARP1. ATR inhibition causes a unique type of replication stress through upregulation of CDK1/2 activity, which increases origin firing and suppresses RRM2 ^5,14–17,20^. The increase in CDK1/2 activity mediated by ATRi results in the slowing down of the replication forks, the uncoupling of helicases and polymerases, and an accumulation of ssDNA ^1,3^. Cell treatment with CDK1/2 inhibitors (PHA-793887 [PHA] or Roscovitine [Rosc]) fully suppressed PARP1 MAR/PARylation and PARP1 trapping after both ATRi and Chk1i treatments **(Figures 3A-B and Supplementary Figures 2A-B)**, suggesting that the surge in origin firing caused by ATRi or Chki is responsible for PARP1 hyperactivation. Similarly, cells treated with HU or aphidicolin (APH) to block ongoing replication forks and thereby inhibiting the progression of the new origins of replication failed to activate PARP1 upon treatment with ATRi or Chk1i **(Figures 3C-D and Supplementary Figure 2C)**. These results demonstrate that PARP1 hyperactivation in replicating cells is triggered by a specific type of replication stress resulting from ATR inhibition-mediated new origin firings.

**Figure 3:**
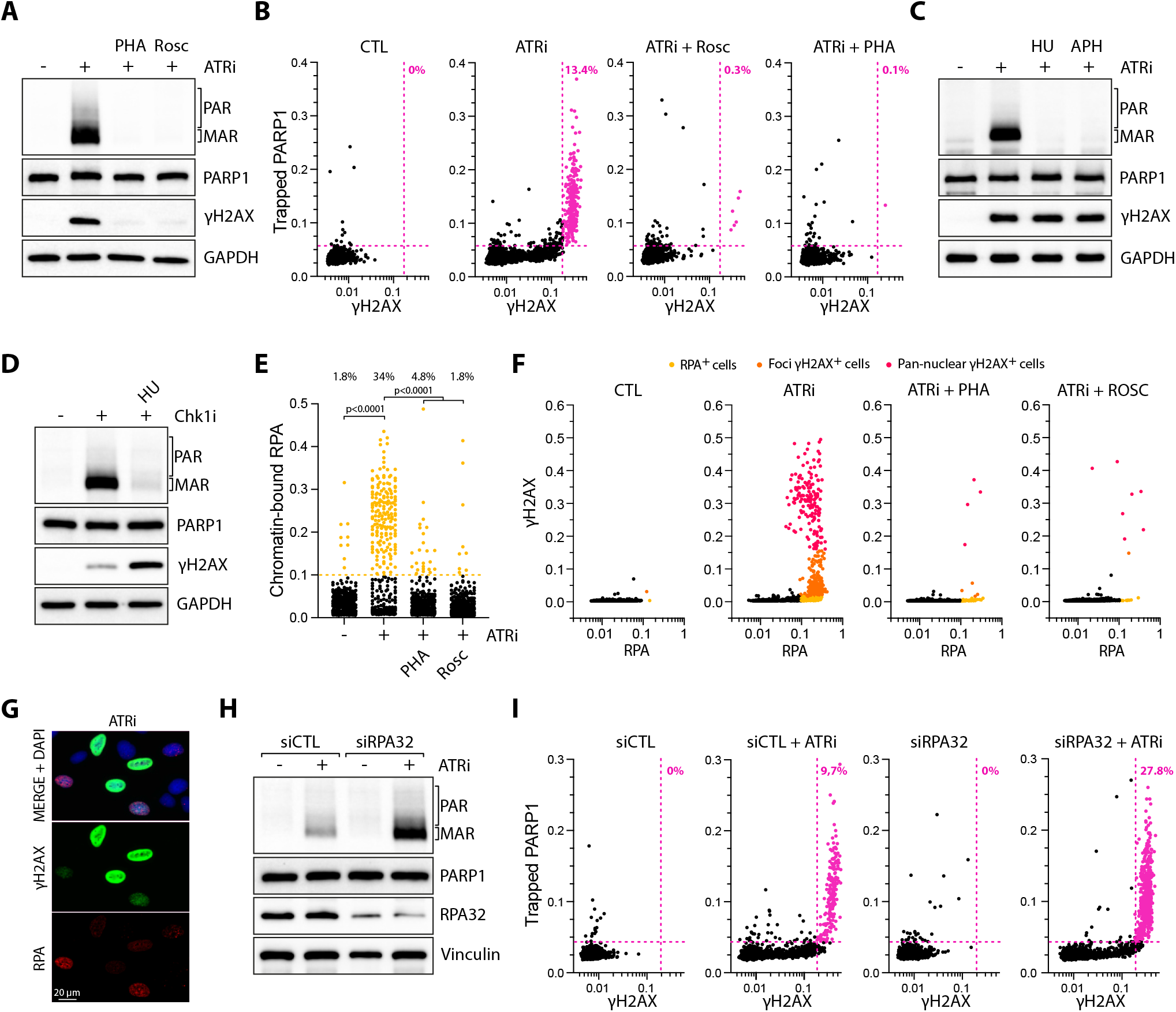
Unprotected ssDNA mediated by aberrant origin firing leads to PARP1 hyperactivation. **A-B**. U2OS cells were treated with ATRi (1 μM) ± Roscovitine (12.5 μM) or PHA-793887 (3 μM) for 4h and analyzed by western blot **(A)** or by with immunofluorescence (number of cells, n=2,000) **(B)** with indicated antibodies. Cells were pre-extracted to remove soluble proteins before performing immunofluorescence. **C-D**. U2OS cells were treated with the indicated drugs for 4h and analyzed by western blot with antibodies against poly/mono ADP-ribose, PARP1, γH2AX, and GAPDH. **E**. Quantification of chromatin-bound RPA by immunofluorescence of 500 U2OS cells treated with ATRi (1 μM) ± PHA-793887 (3 μM) or Roscovitine (12.5 μM) for 4h. Top: Percentage of RPA-positive cells. **F**. Immunofluorescence quantification of γH2AX and chromatin-bound RPA levels in U2OS cells treated with ATRi (1 μM) ± PHA-793887 (3 μM) or Roscovitine (12.5 μM) for 4h (number of cells, n=2,000). Cells were color-coded as follows: yellow for RPA-positive cells only, orange for RPA-positive cells with low γH2AX intensity, and red for RPA-positive cells with high γH2AX intensity. **G**. Representative immunofluorescence picture of U2OS cells treated with ATRi for 4h and stained for γH2AX and RPA. Scale bar: 20 μm. **H-I**. U2OS cells were transfected with siRNA CTL or against RPA32 (0.01 nM) for 40h and subsequently treated with ATRi (1 μM) for 4h. Cells were then analyzed by western blot with indicated antibodies **(H)** or by immunofluorescence against PARP1 and γH2AX (number of cells, n=2,000) (I). Colored dots and percentages indicate cells positive for γH2AX and PARP1. All P-values were calculated with a two-tailed Student t-test.

The surge in new origin firing in cells following inhibition of the ATR pathway leads to increased levels of ssDNA associated with DNA replication forks ^5,20^. In unstressed cells, RPA protects ssDNA formed at replication forks during DNA replication. Following ATR inhibition, the increase in ssDNA is directly linked to elevated RPA intensity in replicating cells **(Figure 3E)** ^5,20^. However, if too many origins are fired simultaneously, there is not enough RPA molecules in cells to adequately cover all of the ssDNA at replication forks, leaving a significant fraction of ssDNA unprotected and vulnerable to breakage, resulting in fork collapse and γH2AX activation **(Figures 3A-B)**^19,20^. Consistent with previous studies ^5,20^, RPA-positive cells gradually become positive for γH2AX following ATRi treatment **(Supplementary Figure 2D)**, illustrating how RPA exhaustion results in ssDNA breakage at replication forks. Inhibition of CDK1/2 prevents both the increase of RPA and γH2AX levels in ATRi-treated cells **(Figures 3E-G)**, further confirming that the surge in origin firing caused by ATR inhibition leads to ssDNA formation followed by replication fork collapse and DSB formation 20. We next asked whether RPA protects cells from PARP1 hyperactivation. We partially knocked down RPA32 to decrease the levels of available RPA in cells, impairing RPA’s ability to protect ssDNA after ATRi treatment without affecting cell replication **(Supplementary Figure 2E)** ^20^. We found that the decrease in RPA levels further enhanced both PARP1 MAR/PARylation levels and PARP1 trapping **(Figures 3H-I)**. These results demonstrate that unprotected ssDNA at replication forks, due to RPA exhaustion in cells leads to PARP1 hyperactivation.

### APOBEC3B triggers PARP1 hyperactivation at replication forks

Given that APOBEC-induced mutations are associated with the lagging strand of the replication forks ^29–31^, we tested whether the hyperactivation of PARP1 after ATRi was a result of APOBEC activity targeting unprotected ssDNA at replication forks. We knocked down or knocked out (KO) A3B, which is highly expressed in U2OS cells ^76^, and monitored PARP1 activation following ATRi treatment. In the absence of A3B, both PARP1 MAR/PARylation levels and PARP1 trapping were completely abrogated following ATRi or Chk1i treatment **(Figures 4A-C and Supplementary Figures 3A-B)**, demonstrating that A3B is essential for PARP1 hyperactivation. Importantly, cells knocked down for A3B and complemented with A3B wild-type restored MAR/PARylation levels after ATRi **(Figure 4D)**, further supporting that A3B is the key enzyme causing PARP1 hyperactivation in response to ATR inhibition. Likewise, A3B knockdown suppressed MAR/PARylation levels in TOV21G and T98G cells, two other cell lines expressing high levels of A3B **(Supplementary Figures 3C-D)**. Next, we selected the SKBR3 cell line, which lacks the A3B gene ^77^, and the MCF10A cell line, which expresses a very low level of A3B ^37^. As expected, both cell lines failed to activate PARP1 in response to ATRi treatment, in contrast to U2OS cells that express A3B **(Figure 4E and Supplementary Figure 3E)**. Nevertheless, A3B overexpression further enhances PARP1 activation in response to ATRi treatment, while the A3B catalytic dead mutant (A3B^E255Q^) did not **(Figure 4F)**, demonstrating that A3B deaminase activity is required for PARP1 activation upon ATRi. Like A3B, A3A wild-type overexpression but not the A3A catalytic dead mutant (A3^AE72A^) promotes PARP1 activation **(Figures 4G-H)**. Finally, we treated cells with MNNG and monitored PARP1 activation in wild-type and A3B KO cells. Although MNNG strongly induced PARP1 activity, both PARP1 MAR/PARylation and trapping levels were not impacted by the absence of A3B in cells **(Figures 4I-J)**. In addition, MNNG treatment induced MAR/PARylation in both SKBR3 and MCF10A cells at levels comparable to those observed in U2OS cells **(Supplementary Figure 3F)**, establishing that the absence of PARP1 activation upon ATRi treatment in these two cell lines was not due to a general defect in PARP1. Taken together, our findings demonstrate that A3B stimulates PARP1 by generating a specific type of DNA lesion at replication forks in response to ATR inhibition, while MNNG generates DNA lesions recognized by PARP1 independently of A3B deaminase activity. Therefore, ATR plays a critical role in protecting replication forks from A3B-induced PARP1 hyperactivation.

**Figure 4:**
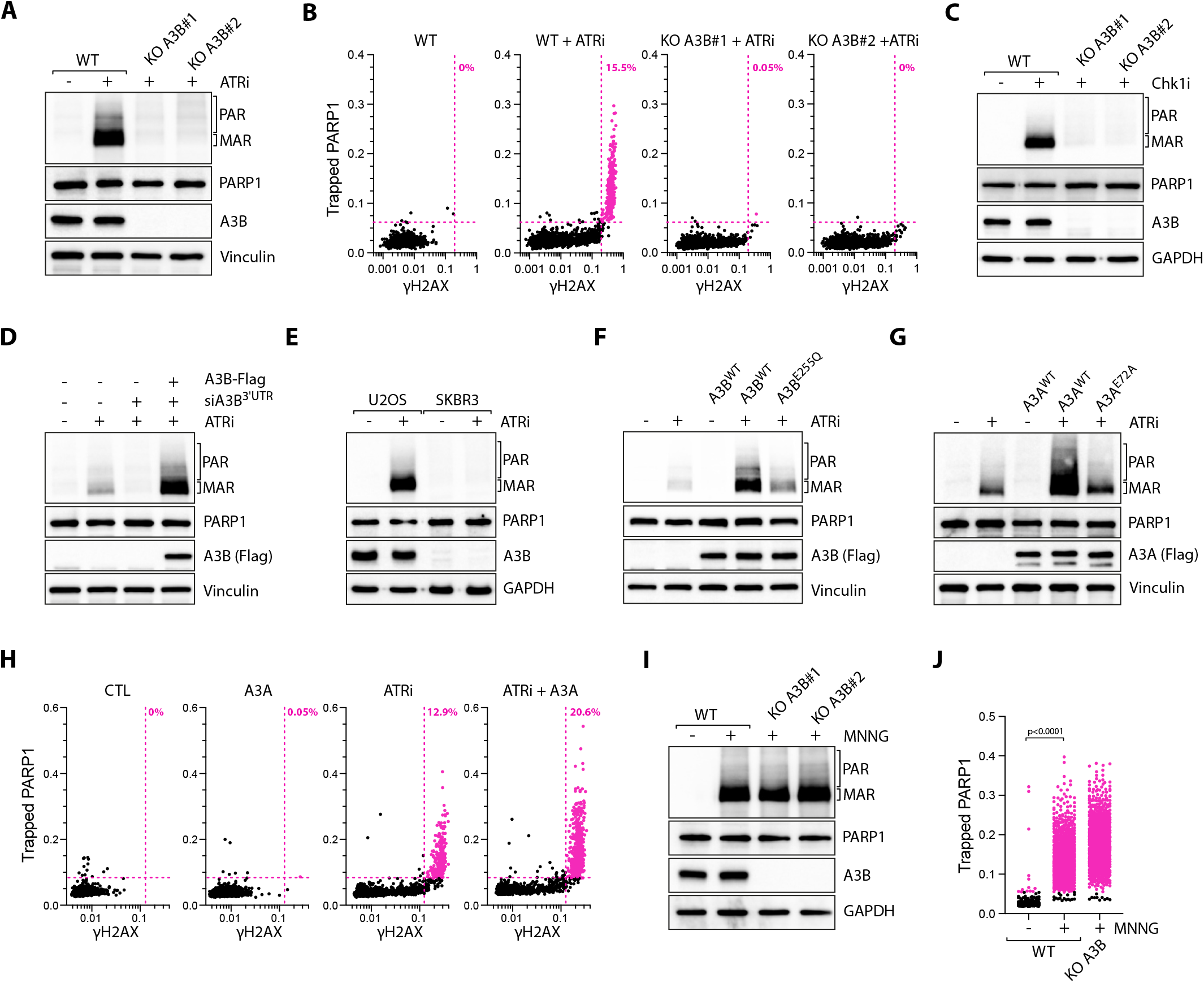
APOBEC3B and APOBEC3A promote PARP1 hyperactivation at replication forks. **A-B**. U2OS WT or A3B KO cells were treated with ATRi (1 μM) for 4h and analyzed by western blot **(A)** or by immunofluorescence (number of cells, n=2,000) **(B)** with indicated antibodies. Colored dots and percentage indicate cells positive for γH2AX and PARP1. **C**. The levels of MAR/PAR were monitored by western blot in U2OS WT or A3B KO cells treated with Chk1i (1 μM) for 4h. **D**. U2OS-A3B-flag cells ± DOX were transfected with an siRNA targeting the 3′UTR of endogenous A3B mRNA or a control siRNA (siCTL) for 40 h and subsequently were treated with ATRi (1 μM) for 4h. The levels of poly/mono ADP-ribose, PARP1, Flag (A3B), and Vinculin were detected by western blot. **E**. U2OS or SKBR3 cells were treated with ATRi (1 μM) for 4h and analyzed by western blot with antibodies against poly/mono ADP-ribose, PARP1, A3B, and GAPDH. **F-G**. U2OS expressing indicated constructs were treated with ATRi (1 μM) for 4h. The levels of poly/mono ADP-ribose, PARP1, Flag (A3B or A3A), and /or Vinculin were analyzed by western blot. **H**. Quantification by immunofluorescence of chromatin-bound PARP1 and γH2AX levels of 2,000 U2OS-A3A-flag cells ± DOX treated with ATRi (1 μM) for 4h. Colored dots and percentage indicate cells positive for γH2AX and PARP1. **I**. U2OS WT or A3B KO cells were treated with DMSO and MNNG (10 μM) for 15 min and analyzed by western blot with antibodies against the indicated proteins. **J**. PARP1 trapping was monitored by immunofluorescence in 2,000 U2OS WT or A3B KO cells treated with MNNG (10 μM) for 1h. Colored dots indicate cells positive for PARP1. All P-values were calculated with a two-tailed Student t-test.

### APOBEC3B causes replication fork collapse upon ATR inhibition

To further investigate the role of A3B activity in promoting PARP1 at replication forks, we first monitored chromatin-bound RPA following ATR inhibition. We found that A3B KO cells did not affect the increased levels of RPA intensity in ATRi-treated cells **(Figure 5A)**, suggesting that A3B acts after the formation of ssDNA caused by the surge of origin firing upon ATRi treatment. We then monitored the levels of γH2AX and chromatin-bound RPA. While the absence of A3B did not affect RPA levels, it significantly suppressed γH2AX activation **(Figure 5B)**. These results demonstrate that A3B drives the breakage of unprotected ssDNA and the formation of DSBs resulting from RPA exhaustion caused by the surge of new origin firing. Previously, we reported that γH2AX pan-nuclear activation following ATR inhibition was mediated by DNA-PK kinase ^5^. Like PARP1 MAR/PARylation levels, DNAPKcs phosphorylation was suppressed after ATRi or Chk1i treatment in A3B KO cells **(Figures 5C-D)**. Although DNA-PK inhibition reduced γH2AX levels, it did not affect PARP1 MAR/PARylation levels following ATRi treatment **(Figure 5E)**. Inversely, PARPi had no impact on DNA-PKcs phosphorylation levels **(Supplementary Figure 3G)**. Thus, DNA-PK and PARP1 are independently activated at broken replication forks mediated by A3B after ATR inhibition. Finally, we monitored how A3B activity affects XRCC1 recruitment to replication forks upon ATRi treatment. In the absence of A3B, XRCC1 localization to replication forks was completely abrogated **(Figure 5F)**, suggesting that A3B-mediated PARP1 activation is critical for the recruitment of XRCC1 to collapsed replication forks. Taken together, these results demonstrate that A3B directly targets unprotected ssDNA at replication forks, causing replication fork breakage, activating PARP1 and DNAPKcs, and subsequently leading to the recruitment of repair factors.

**Figure 5:**
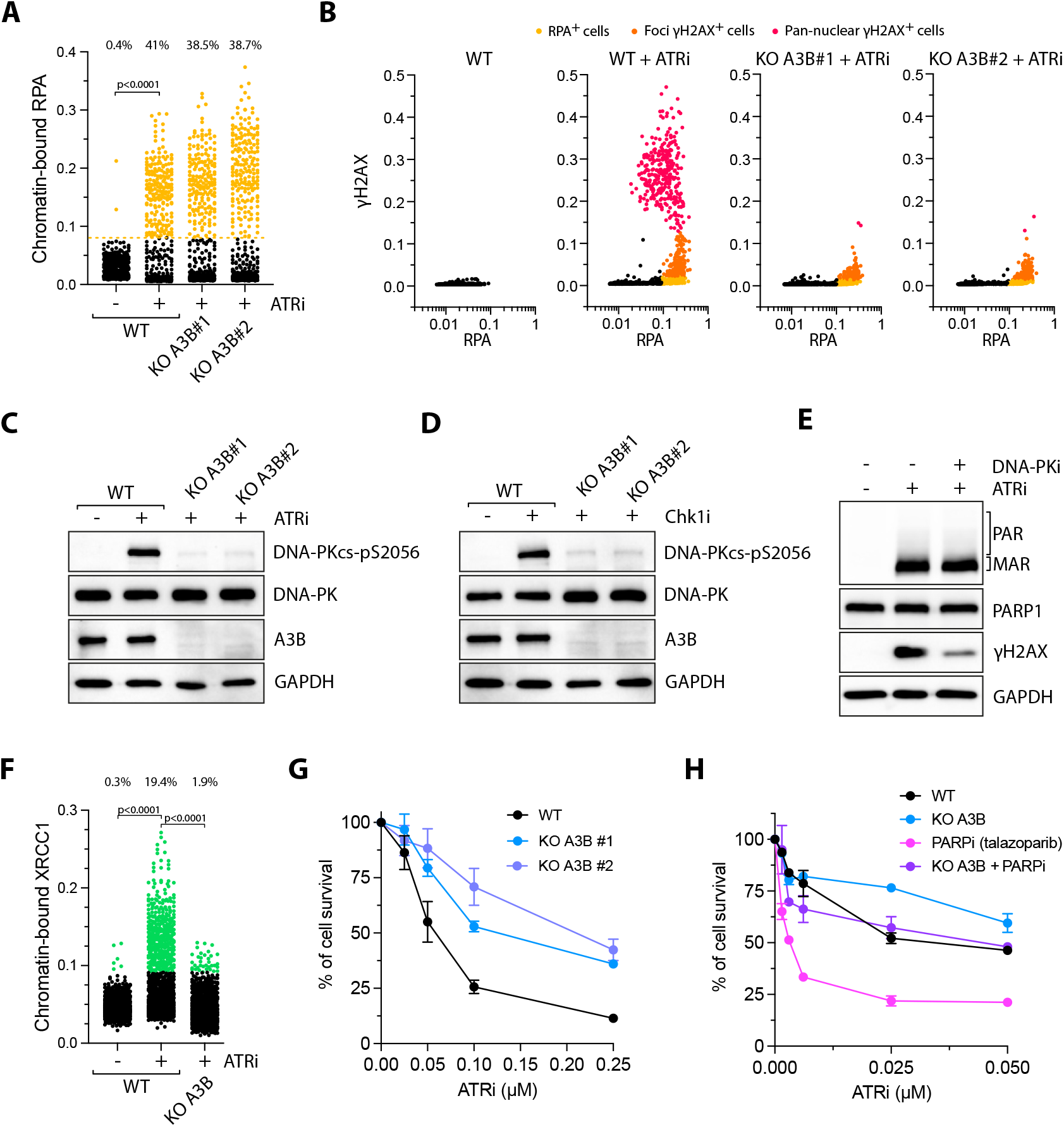
APOBEC3B promotes ATR-mediated replication fork collapse. **A**. Quantification of chromatin-bound RPA by immunofluorescence of 500 U2OS WT or A3B KO cells treated with ATRi (1 μM) for 4h. Top: Percentage of RPA-positive cells. **B**. U2OS WT or A3B KO cells were treated with ATRi (1 μM) for 4h and analyzed by immunofluorescence with RPA and γH2AX antibodies (number of cells, n=2,000). Cells were color-coded as follows: yellow for RPA-positive cells only, orange for RPA-positive cells with low γH2AX intensity, and red for RPA-positive cells with high γH2AX intensity. **C-D**. U2OS WT or A3B KO cells were treated with ATRi (1 μM) **(C)** or Chk1i (1 μM) **(D)** for 4h and analyzed by western blot with antibodies against DNA-PKcs-pS2056, DNA-PKcs, A3B, and GAPDH. **E**. The levels of poly/mono ADP-ribose, PARP1, γH2AX, and GAPDH were analyzed by western blot following treatment of U2OS cells with ATRi (1 μM; 4h) or DNA-PKi (5 μM; 4h). **F**. Chromatin-bound XRCC1 intensity quantification by immunofluorescence of 2,000 U2OS WT or A3B KO cells treated with ATRi (1 μM) for 4h. Top: Percentage of XRCC1-positive cells. **G**. Cell survival of U2OS WT or A3B KO cells treated with increasing concentrations of ATRi for 24 h and then cultured in inhibitor-free media for 48 h. Mean values ± SEM. Number of biological replicates, n = 3. **H**. Cell survival of U2OS WT or A3B KO cells treated with increasing concentration of ATRi ± PARPi (1 μM talazoparib) for 24 h and then cultured in inhibitor-free media for 48 h. Mean values ± SEM. Number of biological replicates, n = 3. All P-values were calculated with a two-tailed Student t-test.

### Endogenous APOBEC3B does not promote ssDNA gaps at ongoing replication forks

One consequence of ectopic overexpression of A3A in cells is the formation of toxic ssDNA gaps behind replication forks through PrimPol-mediated repriming ^55,78^. To test whether the expression of endogenous A3B causes ssDNA gaps at replication forks, we labeled nascent DNA with the thymidine analogs 5-chloro-2′-deoxyuridine (CldU) and 5-iodo-2′-deoxyuridine (IdU) and performed the DNA fiber assay in the presence or absence of S1 nuclease, which specifically cleaves ssDNA gaps ^79^. We measured the length of CldU-labeled replication tracts in U2OS wild-type cells and cells depleted of A3B. We found no change in the CIdU tract length after digestion with the S1 nuclease **(Supplementary Figure 4A)**, indicating that A3B does not induce a significant amount of ssDNA gaps. Thus, the endogenous expression of A3B in U2OS cells does not induce genotoxic ssDNA gaps at ongoing replication forks, potentially explaining why cancer cells can tolerate constitutively high expression levels of A3B but not A3A. Of note, U2OS cells are among the cancer cells expressing the highest levels of A3B ^76^. Furthermore, after ATRi treatment, the knockdown of PrimPol did not impact PARP1 MAR/PARylation levels (**Supplementary Figures 4B-C)**, indicating that PrimPol-generated ssDNA gaps are not required for PARP1 hyperactivation after ATR inhibition. Instead, we propose that ATR inhibition generates substrates for A3B at replication forks, allowing A3B to induce specific types of toxic DNA damage that trigger PARP1 hyperactivation.

### APOBEC3B-mediated PARP trapping confers cell hypersensitivity to ATR inhibitor

Although ectopic overexpression of A3A or A3B confer cell sensitivity to ATR inhibitors ^46,48^, it is still unknown whether endogenous A3B levels are sufficiently high to trigger cell death upon ATRi treatment. To address this, we treated A3B KO cell lines with increasing concentrations of ATRi and observed significant resistance to ATR inhibition when compared to wild-type cells **(Figure 5G)**. Similar resistance was obtained in A3B KO cells treated with Chk1i **(Supplementary Figure 4D)**. These results reveal that endogenous expression of A3B is sufficient to confer cell sensitivity to ATR or Chk1 inhibition. In addition, if A3B is the key factor responsible for fork collapse and PARP1 hyperactivation upon ATR inhibitor treatment, one would predict that PARP inhibitors should render cells more sensitive to ATRi in an A3B-dependent manner. Indeed, PARP inhibitors combined with ATRi enhance synergistically cell death and A3B depletion strongly reduced cell sensitivity to combined ATRi and PARPi treatment **(Figure 5H and Supplementary Figure 4E)**. Importantly, cell sensitivity to ATRi in combination with PARPi was independent of PrimPol **(Supplementary Figure 4F)**, further demonstrating that PARPi-mediated ssDNA gaps were not responsible for the enhanced cell death upon ATRi treatment. Together, these results further support the idea that A3B-induced toxic PARP1 trapping in response to ATRi drives cell sensitivity to ATR inhibition, leading to a context of synthetic lethality when combined with PARP inhibitors.

### Cleavage of abasic sites by APE1 triggers PARP1 hyperactivation

We next investigated the downstream mechanisms by which APOBEC promotes fork collapse and activates PARP1 upon ATR inhibition. We first knocked down or knocked out UNG2, which is the main glycosylase removing uracils from DNA and leading to the formation of abasic sites (or apurinic/ apyrimidinic [AP] site). Moreover, UNG2 is responsible for APOBEC-mediated mutational signature SBS13 corresponding to C to T and C to G mutations in cancer genomes ^24,35^. In the absence of UNG2, the increased levels of RPA associated with replication forks remained unchanged after ATR inhibition **(Figure 6A)**. However, MAR/PARylation levels and PARP1 trapping were completely abolished **(Figures 6B-C and Supplementary Figure 5A)**, similar to the results obtained with A3B KO cells **(Figures 4A-B and 5A)**. These findings suggest that the formation of abasic sites by UNG2 is a required step for PARP1 activation upon ATR inhibition and occurs after the formation of unprotected ssDNA at replication forks. We then investigated how A3B-mediated abasic site formation on unprotected ssDNA contributes to replication fork collapse. We knocked down or knocked out apurinic/apyrimidinic endonuclease 1 (APE1), which cleaves abasic sites in both ssDNA and double-stranded DNA (dsDNA) ^80,81^. Like in A3B and UNG2 KO cells, RPA chromatin-bound levels were not impacted in APE1 KO compared to wild-type cells following ATR inhibition **(Figure 6D)**, demonstrating that the formation of ssDNA was not affected by the absence of APE1. However, both PARP1 MAR/PARylation levels and PARP1 trapping were suppressed in APE1 KO cells treated with ATR inhibition **(Figures 6E-F and Supplementary Figure 5B)**, demonstrating that APE1mediated cleavage of abasic sites at replication forks induces PARP1 hyperactivation. Furthermore, DNA-PKcs phosphorylation and γH2AX levels were suppressed in both UNG2 and APE1 KO cells **(Figure 6G and Supplementary Figures 5C-D)**, further supporting that APE1-mediated cleavage of UNG2-induced abasic sites drives replication fork collapse. Finally, we asked whether UNG2 and APE1 also impact XRCC1 recruitment to collapsed replication forks. Similar to A3B KO cells, both UNG2 and APE1 KO cells showed reduced chromatin association of XRCC1 **(Figure 6H)**, confirming that PARP1 activation at collapse forks facilitates the recruitment of repair factors. Collectively, these results demonstrate that following A3B-mediated deamination of ssDNA, UNG2 and APE1 are crucial for removing deaminated cytosines and cleaving abasic sites, leading to fork collapse, PARP1 activation, and repair factor recruitment.

**Figure 6:**
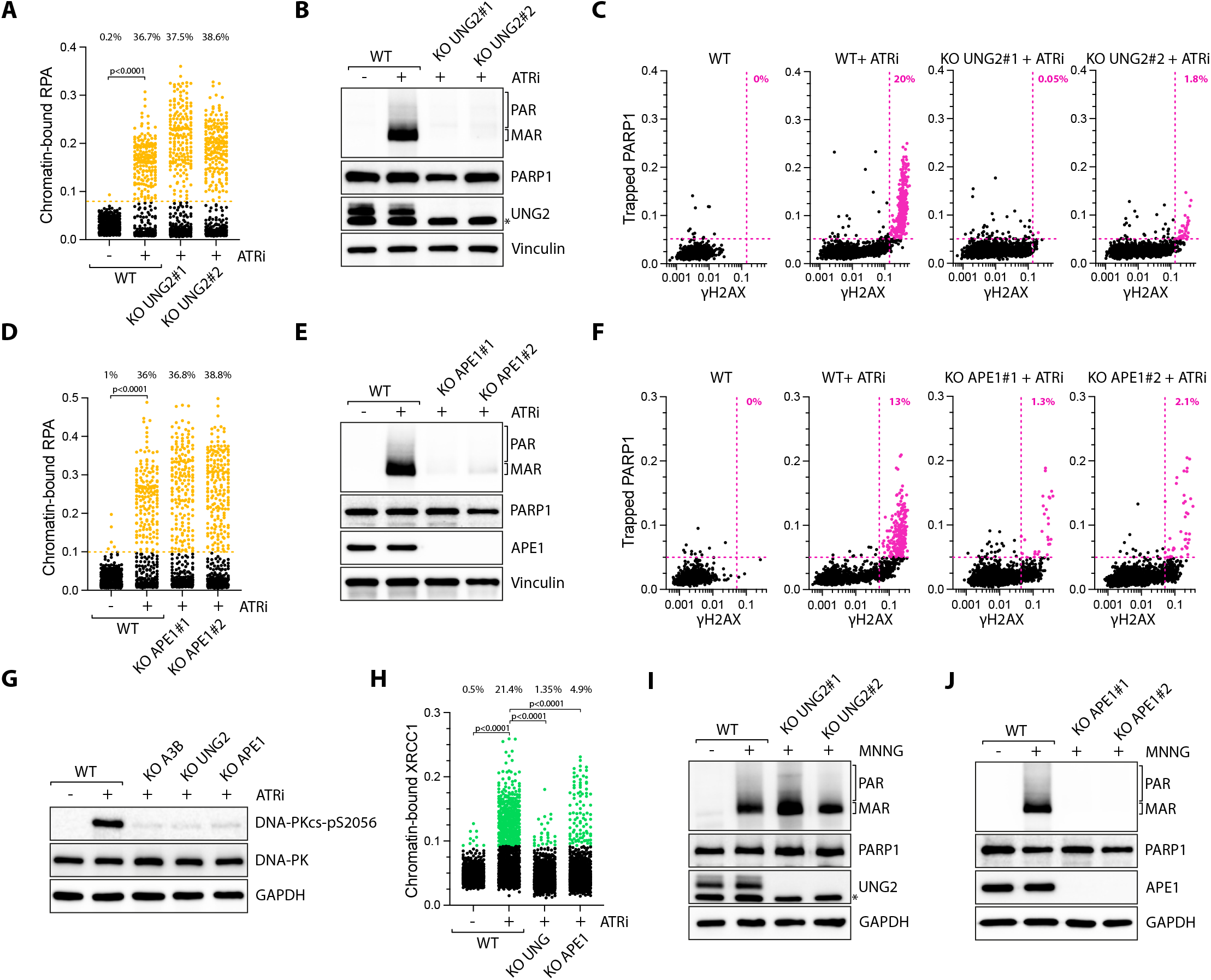
Abasic site cleavage by APE1 induces PARP1 hyperactivation. **A**. Chromatin-bound RPA intensity quantification by immunofluorescence of 2,000 U2OS WT or UNG2 KO cells treated with ATRi (1 μM) for 4h. Top: Percentage of RPA-positive cells. **B-C**. U2OS WT or UNG2 KO cells were treated with ATRi (1 μM) for 4h and analyzed by western blot **(B)** or by with immunofluorescence (number of cells, n=2,000) **(C)** with indicated antibodies. Colored dots and percentages indicate cells positive for γH2AX and PARP1. **D**. Chromatin-bound RPA intensity quantification by immunofluorescence of 2,000 U2OS WT or APE1 KO cells treated with ATRi (1 μM) for 4h. Top: Percentage of RPA-positive cells. **E-F**. U2OS WT or APE1 KO cells were treated with ATRi (1 μM) for 4h and analyzed by western blot **(E)** or by immunofluorescence (number of cells, n=2,000) **(F)** with indicated antibodies. Colored dots and percentages indicate cells positive for γH2AX and PARP1. **G**. The levels of DNA-PK-pS2056, DNA-PK, and GAPDH were analyzed by western blotting following ATRi (1 μM; 4h) in the indicated U2OS cell lines. **H**. U2OS WT, UNG2 KO, or APE1 KO cells were treated with ATRi (1 μM) for 4h and analyzed by immunofluorescence with XRCC1 and γH2AX antibodies (number of cells, n=2,000). Top: Percentage of XRCC1-positive cells. **I-J**. U2OS WT, UNG2 KO or APE1 KO cells were treated with MNNG (10 μM) for 15 min and analyzed by western blot with antibodies against poly/mono ADP-ribose, PARP1, UNG, APE1, and GAPDH. All P-values were calculated with a two-tailed Student t-test.

While APE1 is the key enzyme driving PARP1 hyperactivation resulting from A3B activity at replication forks, other nucleases may also participate in fork cleavage after ATRi. Previous studies reported MUS81-dependent DNA damage after ATRi and Chk1i ^5,82–85^. As expected, MUS81 knockdown significantly reduced γH2AX levels, but PARP1 MAR/ PARylation levels remained unaffected in the absence of MUS81**(Supplementary Figures 6A-B)**, indicating that MUS81-mediated fork cleavage generates distinct types of DNA break end-products that do not trigger PARP1 hyperactivation. A recent study revealed RAD51 as a crucial factor that directly binds to abasic sites, preventing their cleavage and the formation of DNA breaks ^86^. However, neither the knockdown of RAD51 nor BRCA2 (an essential factor for RAD51 recruitment) significantly affected PARP1 activation after ATRi treatment **(Supplementary Figures 6C-D)**, suggesting that these repair factors are not involved in A3B-mediated PARP1 activation in response to ATR inhibition. ATR is also known to promote RAD51 recruitment to DNA breaks through the regulation of the BRCA1-PALB2-BRCA2 complex, which is essential for RAD51 loading on ssDNA ^6,87^. Thus, ATR inhibition may not only facilitate the formation of abasic sites by increasing the level of ssDNA targeted by A3B, but also by preventing RAD51 protection mechanisms that suppress abasic site cleavage.

### APE1-induced PARP1 hyperactivation occurs regardless of how abasic sites are generated

To better understand the global mechanism driving PARP1 hyperactivation in cells, we examined the specific roles of UNG2 and APE1 in response to other DNA-damaging agents known to trigger PARP1’s hyperactivation. We first treated UNG2 or APE1 KO cells with MNNG. MNNG causes DNA methyl adducts, inducing the formation of abasic sites directly through the labilization of the N-glycosidic bonds between bases and sugars independently of replication ^88^. Unlike ATR inhibition, MNNG treatment induced MAR/PARylation levels in both wild-type and UNG2 KO cells **(Figure 6I)**. This result is consistent with MNNG’s ability to directly generate abasic sites, bypassing the need for both UNG2 and A3B. In contrast, APE1 KO completely abrogated PARP1-associated MAR/PARylation **(Figure 6J)**, demonstrating that abasic site cleavage is the critical step for PARP1 hyperactivation. We further validated these results by treating cells with MMS (methyl methanesulfonate) and H_2_O_2_, well-known chemical compounds that induce abasic sites and hyperactivate PARP1^72,89^. Like ATRi and MNNG, APE1 KO cells failed to activate PARP1 after treatment with MMS or H_2_O_2_, while UNG2 KO cells retained high MAR/PARylation levels **(Supplementary Figures 7A-D)**. However, cells treated with ETP, CPT, or HU in the presence of PARGi triggered PARP1 MAR/PARylation in an APE1-independent manner **(Supplementary Figures 7E)**, further implying that APE1-mediated PARP1 hyperactivation only occurs in response to DNA-damaging agents inducing abasic sites in DNA. We next monitored PARP1 trapping and γH2AX levels upon MNNG treatment. Consistently with the MAR/PARylation level results, MNNG-induced PARP1 trapping was absent in APE1 KO cells **(Supplementary Figure 7F)**. However, unlike ATRi treatment, where APE1 was responsible for γH2AX induction **(Figures 6E-F)**, the absence of APE1 did not reduce γH2AX levels **(Supplementary Figure 7F)**, suggesting that PARP1 hyperactivation occurs independently of the formation of DNA breaks caused by MNNG treatment. Therefore, PARP1 trapping on DNA does not directly correlate with the overall levels of DNA breaks in cells but is instead determined by the presence of a specific type of breaks generated by APE1. These findings collectively show that the cleavage of abasic sites by APE1 is the critical step in promoting PARP1 hyperactivation in cells, independent of the mechanisms that generate the abasic sites, DNA break levels, or whether the abasic sites occur at replication forks or within double-stranded DNA.

**Figure 7:**
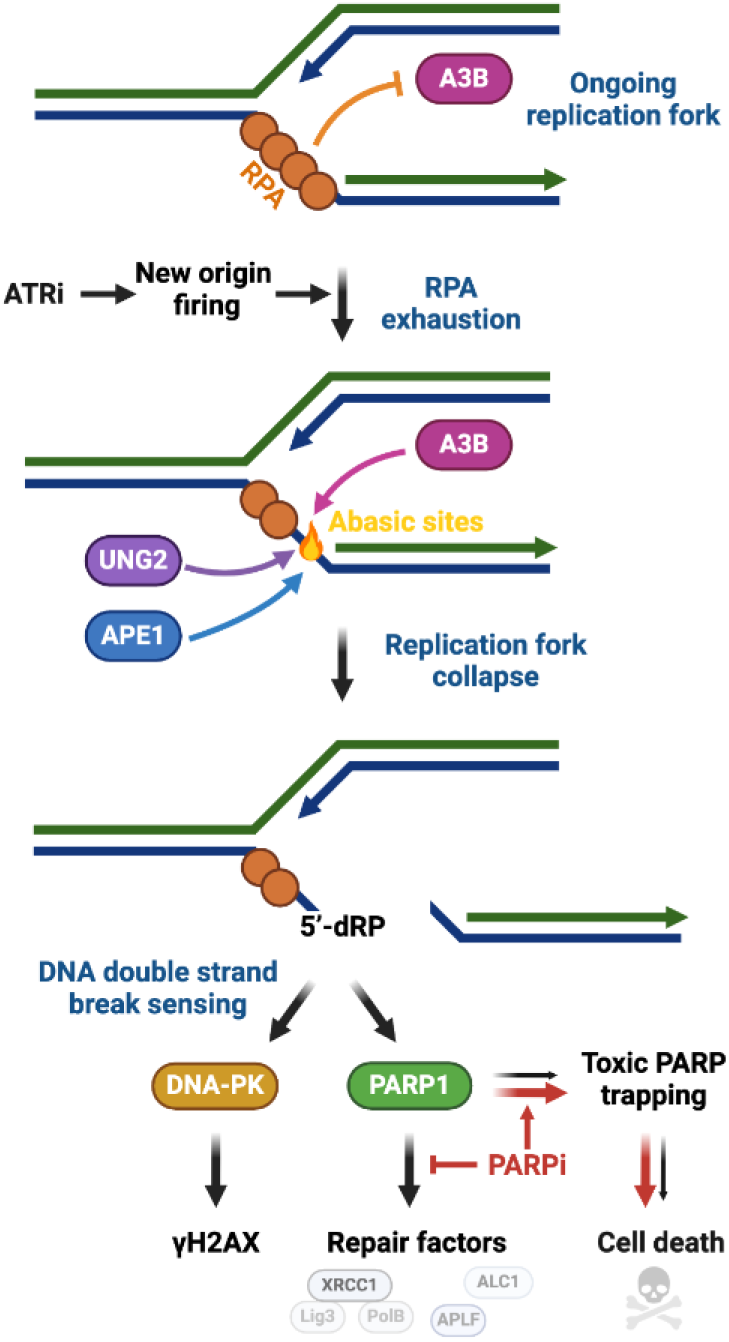
Working Model. Following ATR inhibition, the surge of origin firing leads to RPA exhaustion, rendering ssDNA at replication forks unprotected from nuclear attacks. A3B promotes the deamination of cytosine on unprotected ssDNA, which are removed by UNG2 to form abasic sites. Abasic sites are subsequently cleaved by APE1, causing forks to collapse and DNA double-strand break formation, which is recognized by DNA-PK and PARP1. PARP1 recruits repair factors such as XRCC1 to replication forks. However, PARP1 trapping increases cellular sensitivity to ATR inhibitors, a vulnerability that is further amplified by PARP inhibitors, which stabilize toxic PARP trapping.

## DISCUSSION

ATR is the master safeguard of genomic integrity during DNA replication by controlling origin firing and promoting the repair of stalled or collapsed replication forks ^1–3^. Inhibition of ATR promotes a surge of new origins firing, leading to a high amount of ssDNA that depletes all available RPA present in cells ^19,20^. RPA exhaustion from the cells leaves ssDNA unprotected and susceptible to breakage, resulting in replication catastrophe ^19,20^. However, the mechanism by which ssDNA breakage occurs remains unclear. Herein, we found that A3B is the key enzyme attacking unprotected ssDNA at replication forks. Mechanistically, we demonstrated that A3B-induced DNA deamination initiates a cascade of reactions involving UNG2 that first removes the uracils to generate abasic sites on ssDNA, followed by APE1 cleavage of the abasic sites, ultimately leading to replication fork collapse, DSBs, PARP1 hyperactivation, and recruitment of DNA repair factors such as XRCC1 **(Figure 7)**. Our findings explained the fundamental role of RPA in preventing replication catastrophe by protecting ssDNA against APOBEC deaminase activity. Moreover, this study provided a mechanistic basis for the synergistic effect between ATR and PARP inhibitors in killing cancer cells expressing high levels of A3B. A3B targets ssDNA generated specifically after acute ATR inhibition, causing toxic PARP1 trapping at replication forks, which is further amplified by PARP inhibitors, establishing a synthetic lethality context **(Figure 7)**.

But why is ATR inhibition so effective at activating PARP1 at replication forks compared to other types of DNA damage that cause fork collapse? Alkylating agents such as MMS and MNNG are also DNA-damaging agents that trigger hyperactivation of PARP1 at a similar level to ATRi but independently of DNA replication. This suggests that treatment with ATR inhibitors and alkylating agents causes specific and common types of DNA lesions in cells, which are particularly efficient at activating PARP1. Indeed, we revealed that PARP1 hyperactivation occurring after ATRi, MNNG, MMS, or H^2^O^2^ strictly depends on the apurinic-apyrimidinic endonuclease APE1 **(Figure 5G and supplementary Figure 5G)**. APE1 cleaves the DNA phosphodiester back bone at the 5’ side of the abasic site in both ssDNA and dsDNA, generating a cleaved DNA strand with a 3’-hydroxyl (3’-OH) and a 5’-deoxyribose phosphate (dRP) termini ^90^, with the latter being the preferred substrate for PARP1^91^. In contrast, other DNA-damaging treatments that cause replication stress and DSBs, such as HU or topoisomerase (TOP) 1 and 2 inhibitors, induce low MAR/PARylation levels detectable only when PARG is absent **(Figure 1E)**. TOP1 and TOP2 inhibitors cause DNA breaks with a 5’or 3’-OH end, respectively, while the second end traps TOP1 or TOP2 ^92^, potentially making poor DNA ends substrates for PARP1. Additionally, MUS81 also cleaves replication forks after ATRi and Chk1i, leading to fork collapse and DSB formation ^5,82–85^. However, our results showed no impact of MUS81 on PARP1 hyperactivation. The lack of effect on PARP1 activation could be attributed to the absence of 5’-dRP group formation at the DNA breaks resulting from MUS81’s DNA cleavage activity. Therefore, we propose that, regardless of the types of DNA-damaging treatments or the levels of DNA breaks, the formation of abasic sites and their subsequent cleavage by APE1 are the key steps necessary to induce PARP1 hyperactivation in damaged cells.

Although A3A/B-induced abasic sites are an essential step for fork cleavage by APE1, other factors or stresses might also promote the formation of abasic sites at replication forks, leading to fork collapse through APE1 activity. The imbalance between dUTP and dTTP in cells is another predominant source of uracil misincorporation into genomic DNA during replication ^93.^ Cell treatment with pemetrexed (PMX) and 5-FU, both thymidylate synthase inhibitors, creates an imbalance in the dUTP/dTTP ratio, leading to elevated levels of genomic uracil and significantly enhancing cell sensitivity to ATR inhibitors ^94^. This suggests that this dUTP/dTTP imbalance could generate similar lesions to those caused by A3A/B activity. However, further studies will be required to determine whether genomic uracils resulting from the UTP/dTTP imbalance are processed through a similar mechanism to uracils induced by A3A/B.

Extensive prior work, including studies from our laboratory, has shown ectopic overexpression of A3A and A3B in cells induces replication stress, DSBs, and cell cycle arrest ^37,46–48,54,56,95,96^. However, whether this accurately reflects the physiological context of A3A and A3B endogenous expression levels remains a topic of debate. For instance, U2OS cells, which express high levels of A3B, replicate without cell cycle arrest or significant levels of DNA damage ^76^. Moreover, we demonstrate that endogenous levels of A3B do not generate ssDNA gaps at ongoing replication forks. This contrasts with ectopic A3A overexpression, which induces both ssDNA gaps and DNA breaks ^37,49,55,78^. A3B’s deaminase activity and DNA binding are much lower than those of A3A due to differences in the structural conformations of their active sites ^44,97^. Consequently, A3B’s DNA binding may be too weak to compete with RPA, whereas A3A can potentially explain the differences between A3A and A3B in causing ssDNA gaps in unstressed cells. Yet, it remains unclear whether endogenous A3A could be expressed at high enough levels in cancer cells to promote ssDNA gaps at replication forks or other types of DNA damage, conferring cell sensitivity to ATRi. This raises concerns about the feasibility of exploiting ATRi in clinics to target cancer cells expressing A3A. Conversely, we showed that endogenous A3B expression, even at the high levels found in cancer cells, cannot counteract RPA’s shielding of ssDNA formed during replication from being targeted by nucleases ^98–100^. A3B deaminates ssDNA only when replication forks encounter obstacles or stress that compromise their protective mechanisms, such as after ATRi. This may explain why cancer cell lines can tolerate high endogenous expression levels of A3B without causing DNA damage or replication defects, which could otherwise be detrimental to tumor growth. Previous models using ectopic A3A/B overexpression suggested that ATR is crucial for repairing DNA damage caused by A3A/B. However, we now propose that cancer cells expressing endogenous A3B are particularly sensitive to ATR inhibitor treatments because ATR inhibition generates the specific substrates targeted by A3B, which, in turn, leads to toxic PARP1 trapping for the cells. This creates a context of synthetic lethality when combined with PARP inhibitors, which further stabilize trapped PARP1 on DNA lesions ^101^. Cells without A3B become resistant to the combination of ATRi and PARPi treatment, suggesting that the A3B expression level is a potential biomarker for determining patients’ tumor sensitivity to these therapies.

## ACKNOWLEDGMENTS

Salary support for P.O. was provided by a California Institute for Regenerative Medicine (CIRM) stem cell biology training grant (TG2-01152) and an EMBO Postdoctoral fellowship (ALTF 213-2023). A.S. and G.S. were supported by the National Institutes of Health Research Supplements to Promote Diversity in Health-Related Research (R37-CA252081-S1;-S2). R.B. was supported by the National Institutes of Health (R37-CA252081) and a Research Scholar Grant (RSG-24-1249960-01-DMC) from the American Cancer Society. A.M.G. was supported by the Department of Defense (CA200867) and Pedal the Cause (FDN-2023-1202). The cartoon in **Figure 7** was generated with BioRender.com.

## AUTHOR CONTRIBUTIONS

P.O., E.B., J.L., A.S., G.S., and B.R.H. performed all the experiments. B.R.H., under the supervision of A.M.G, performed the DNA fibers assay. P.O. and R.B. conceived the study, designed the experiments, and wrote the paper. R.B. oversaw the project, and all the authors contributed to manuscript revisions.

## METHODS

### Plasmids

APOBEC3A and APOBEC3B cDNAs were synthesized by GenScript with a beta-globin intron and a Flag tag in the C-terminus. The plasmids expressing APOBEC3A-Flag or APOBEC3B-Flag were generated by inserting the cDNA into the pInducer20 vector using the Gateway Cloning System (Thermo Fisher Scientific). The catalytically mutants APOBEC3A-E72A and APOBEC3B-E255Q were constructed by site-directed mutagenesis.

### Cell culture

U2OS, HEK-293FT cells were cultured in DMEM supplemented with 10% FBS, 1% L-Glutamine, and 1% penicillin/streptomycin. TOV21G and T98G were cultured in DMEM/F-12 (GlutaMAX) supple mented with 10% FBS and 1% penicillin/streptomycin. SKBR3 cells were cultured in McCoy’s 5A supplemented with 10% FBS and 1% penicillin/ streptomycin. MCF10A cells were cultured in DMEM/F-12 supplemented with 5 % horse serum, 2 ng/mL epidermal growth factor (EGF), 0.5 μg/mL hydrocortisone, 100 ng/mL cholera toxin, 10 μg/mL insulin, and 1 % penicillin/streptomycin. U2OS-derived cell lines were generated by infecting U2OS with lentivirus expressing APOBEC3B, APOBEC3B-E255Q, APOBEC3A, and APOBEC3A-E72A under a doxycycline-inducible promoter (pInducer20) and selected with G418 (750 μg/mL). U2OS-derived cell lines were treated with doxycycline (600 ng/mL) 16 to 24 h before any other indicated treatments.

### Cell treatment

The chemicals and concentrations (if not indicated otherwise) used in this study are listed in **Supplementary Table 1**.

### RNA interference

siRNA transfections were performed by reverse transfection with Lipofectamine RNAiMax (Thermo Fisher Scientific). siRNAs were purchased from Thermo Fisher Scientific (Silencer Select siRNA). Cells were transfected with siRNA for 40h (4-8 nM) before treatment unless indicated. The siRNAs used in this study are listed in **Supplementary Table 2**.

### CRISPR-Cas9 knockout cells

UNG2 and APE1 CRISPR-Cas9 knockout U2OS cell lines were performed by transfection with Lipofectamine CRISPRMAX of TrueGuide Synthetic CRISPR gRNA and TrueCut Cas9 Protein v2 according to the manufacturer’s instructions (Thermo Fisher Scientific). A3B KO-derivative knockout cell lines were generated by transfecting cells with the pSpCas9(BB)-2A-Puro (PX459) plasmid containing gRNAs targeting A3B with FuGENE 6 Transfection Reagent (E2691; Promega). 24h after transfection, cells were selected with puromycin (1 μg/mL). A3B, UNG, and APE1 KO cells were validated by western blot. The gRNAs used in this study are listed in **Supplementary Table 3**.

### Antibodies

The antibodies and dilutions used in this study are listed in **Supplementary Table 4**.

### Immunofluorescence

Cells grown on glass coverslips were incubated in ice-cold pre-extraction buffer (10 mM PIPES pH 6.8, 100 mM NaCl_2_, 300 mM sucrose, 1mM EDTA, and 0.2% Triton X-100) for 5 min on ice, washed twice with PBS 1X and then fixed with paraformaldehyde (3 % paraformaldehyde and 2 % sucrose in 1x PBS) for 20 min. Then, cells were washed twice with 1x PBS and permeabilized with a permeabilization buffer (1x PBS and 0.2 % Triton X-100) for 5 min at room temperature. Subsequently, cells were washed twice with 1x PBS and blocked in PBS-T (1x PBS and 0.05 % Tween-20) containing 2 % BSA and 10 % milk for 1 h. Cells were then incubated with the primary antibody diluted in 1x PBS containing 2 % BSA and 10 % milk at room temperature for 2 h. Coverslips were washed three times with PBS-T for 5 min each wash before 1h incubation with the appropriate secondary antibodies conjugated to fluorophores (Alexa-488 or Cy3). After three 5 min washes with PBS-T, cells were stained with DAPI (5 μg/mL, MilliporeSigma #D9542), and the coverslips were mounted using slow-fade mounting media (Thermo Fisher Scientific, #S36936). When indicated, cells pulse-labeled with EdU (10 μM for 15 min) were labeled using Click-iT EdU Plus EdU Alexa Fluor 594 Imaging Kit (Invitrogen, #C10639) following the fixation and permeabilization steps according to the manufacturer’s protocol. Then, the immunofluorescence protocol was performed as described above. Images were captured using a Leica DMi8 THUNDER microscope.

### Quantitative Image-Based Cytometry

Using a Leica DMi8 THUNDER microscope with a Leica HC PL APO 20x/0,80 objective, 30 to 50 individual images were taken for each sample. The intensity signals of PARP1, γH2AX, XRCC1, and/or RPA32 of individual cells were quantified using the CellProfiler software. The detailed CellProfiler pipeline used to quantify PARP1, γH2AX, XRCC1, and RPA32 intensity levels is described in **Supplementary Methods**. The intensity levels of γH2AX, PARP1, XRCC1, and RPA32 were quantified in DAPI-stained nucleus, and the intensity levels were normalized between the different images by subtracting the background signal from the nuclear intensity signal. Dot blot graphs were generated using GraphPad Prism software.

### Western blotting using the HRP-coupled SpyTag format

Samples were subjected to a standard SDS-PAGE protocol and transferred to PVDF membrane (Sigma, #IPVH00010). The membrane was blocked for 1h at room temperature using a blocking buffer (1x TBS, 0.05 %, Tween-20 and 5% BSA). Then, the membrane was incubated overnight at 4°C in blocking buffer containing 0.05 μg/ml anti-Mono-ADP-Ribose (Bio-Rad, AbD43647 #TZA020), which had been conjugated to BiCatcher2:HRP (Bio-Rad, #TZC002P) according to the manufacturer’s instructions. After incubation, the membrane was washed six times with TBS-T. Protein signals were then detected using SuperSignal West Dura Extended Duration Substrate (Thermo Scientific, #34075) and visualized with a ChemiDoc MP Imaging System (Bio-Rad).

### Cell viability assay

Cells were seeded in 12-well plates at a density of 50,000 cells per well. On the following day, cells were treated with indicated inhibitors. After 24 h of treatment, cells were washed three times with PBS and inhibitor-free media was added. Cell viability was measured 3 days after the treatment using alamarBlue Cell Viability Reagent (ThermoFisher Scientific, #DAL1100) according to manufacturer instructions. Fluorescence levels (excitation 565 nm/emission 590 nm) were measured using a Varioskan LUX Multimode Microplate Reader (ThermoFisher Scientific, #VLBL00GD2).

### Quantitative reverse transcription PCR (RT-qPCR)

Total RNA was extracted from cells using Quick-RNA MiniPrep Kit (Zymo Research, #R1055) according to the manufacturer’s instructions. Following extraction, total RNA was reverse transcribed using the High-Capacity cDNA Reverse Transcription Kit (Thermo Fisher Scientific, #4368813). RT products were analyzed by real-time qPCR using SYBR Green (PowerUp SYBR Green Master Mix, Thermo Fisher Scientific, #A25743) in a QuantStudio 3 Real-Time PCR detection system (Thermo Fisher Scientific). For each sample tested, the levels of indicated mRNA were normalized to the levels of Actin mRNA. The qPCR primers used in this study are listed in **Supplementary Table 5**.

### DNA fiber assay

U2OS cells were first pulse-labeled for 30 min with 20 μM IdU, washed three times with 1X DPBS, and then pulsed with 100 μM CldU for 30 min. After pulse, cells were harvested and collected in icecold DPBS (∼1,500 cells/μL). For the DNA fiber assay with the ssDNA-specific S1 nuclease (S1 Fiber), cells were permeabilized with CSK100 (100 mM NaCl, 10 mM MOPS pH 7, 3 mM MgCl2, 300 mM sucrose and 0.5% Triton X-100 in water) after the CldU pulse for 5 min at room temperature, treated or not with the S1 nuclease (Thermo Fisher Scientific) at 20 U/mL in S1 buffer (30 mM sodium acetate pH 4.6, 10 mM zinc acetate, 5% glycerol, 50 mM NaCl in water) for 30 min at 37 °C, and collected in PBS-0.1% BSA with cell scraper. Nuclei were then pelleted at ∼4700×g for 7 min at 4 °C, then supernatant was aspirated to a final volume of ∼30μL. To spread fibers, 2 μL of cell solution was placed on a charged glass slide, mixed with 6 μL of lysis buffer (200 mM Tris-HCl pH 7.4, 0.5% SDS, 50 mM EDTA), and slides were tilted at 15° to enable gravity to spread DNA fibers. DNA fibers were fixed in a 3:1 solution of methanol and acetic acid for 5 min, denatured in 2.5 M HCl for 1 h, and blocked in pre-warmed 5% BSA at 37 °C for 1 h. IdU and CldU were detected using mouse anti-BrdU (1:20, Invitrogen) and rat anti-BrdU (1:75, Abcam), respectively for 1.5 h at room temperature in a humid chamber followed by anti-mouse Alexa-546 (1:50) and anti-rat Alexa-488 (1:50) secondary antibodies for 1 h at room temperature in a humid chamber. Coverslips were mounted with Prolong Gold Antifade Solution (Invitrogen) and cured overnight at room temperature, protected from light. Fibers were imaged with a 63X oil objective on a Leica DM4 B. Quantification and measurement of fibers was done in ImageJ by blinded analysis.

---

**Supplementary Figure 1:**
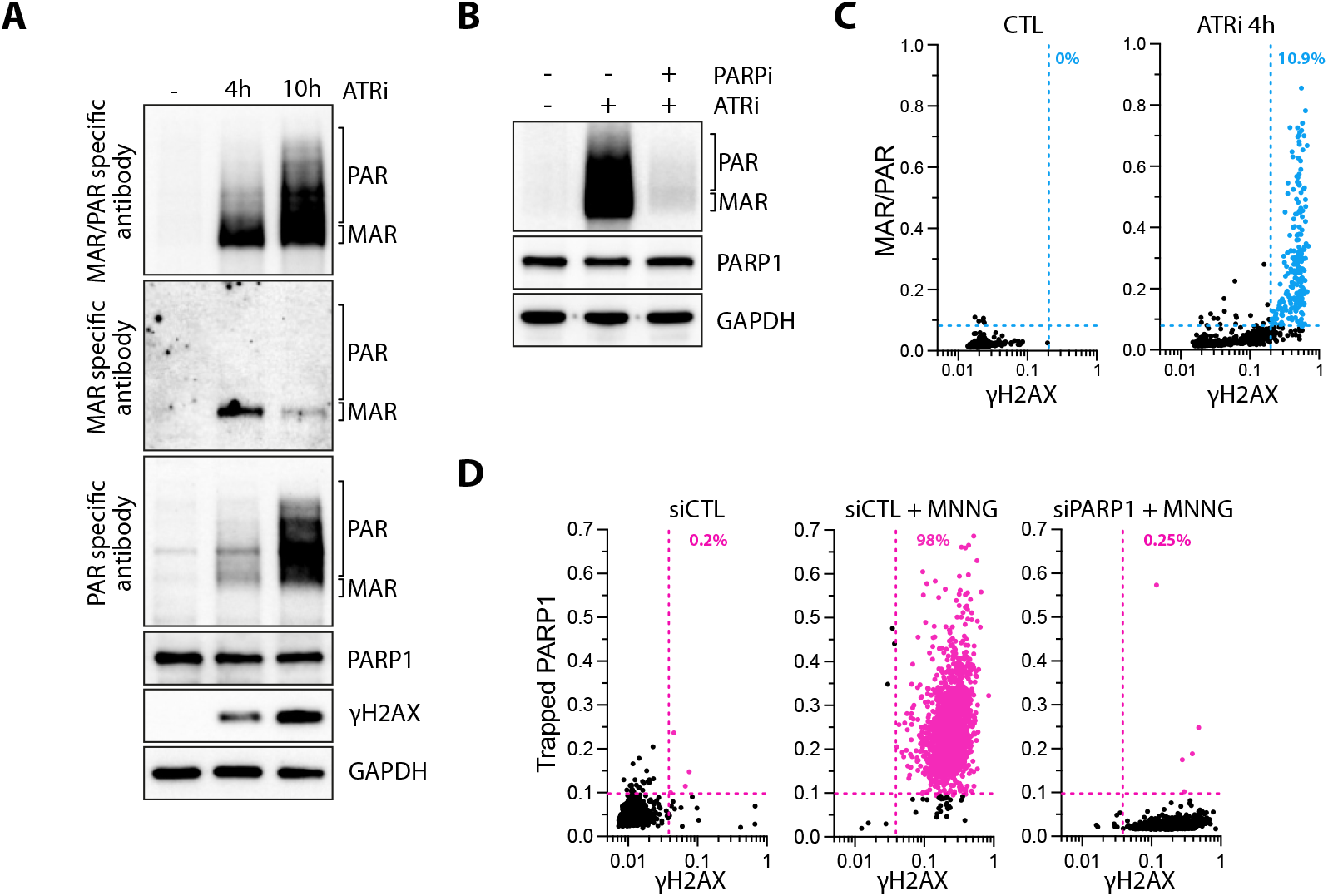
**A**. The levels of MAR/PAR, MAR, or PAR were monitored by western blotting in U2OS cells treated with ATRi (1 μM) for 4h or 10 h. **B**. U2OS cells were treated with ATRi for 4h in the presence or absence of PARPi (10 μM). The levels of poly/mono ADP-ribose, PARP1, and GAPDH were analyzed by western blot. **C**. Quantification of nuclear MAR/PAR and γH2AX in 2,000 U2OS cells treated with ATRi (1 μM) for 4h. Colored dots and percentages indicate cells positive for γH2AX and MAR/PAR. **D**. U2OS cells were transfected with a siRNA control (CTL) or against PARP1 for 40h and subsequently treated with MNNG (10 μM) for 1h and analyzed by immunofluorescence (number of cells, n=2,000) with PARP1 and γH2AX antibodies. Colored dots and percentages indicate cells positive for γH2AX and PARP1.

**Supplementary Figure 2:**
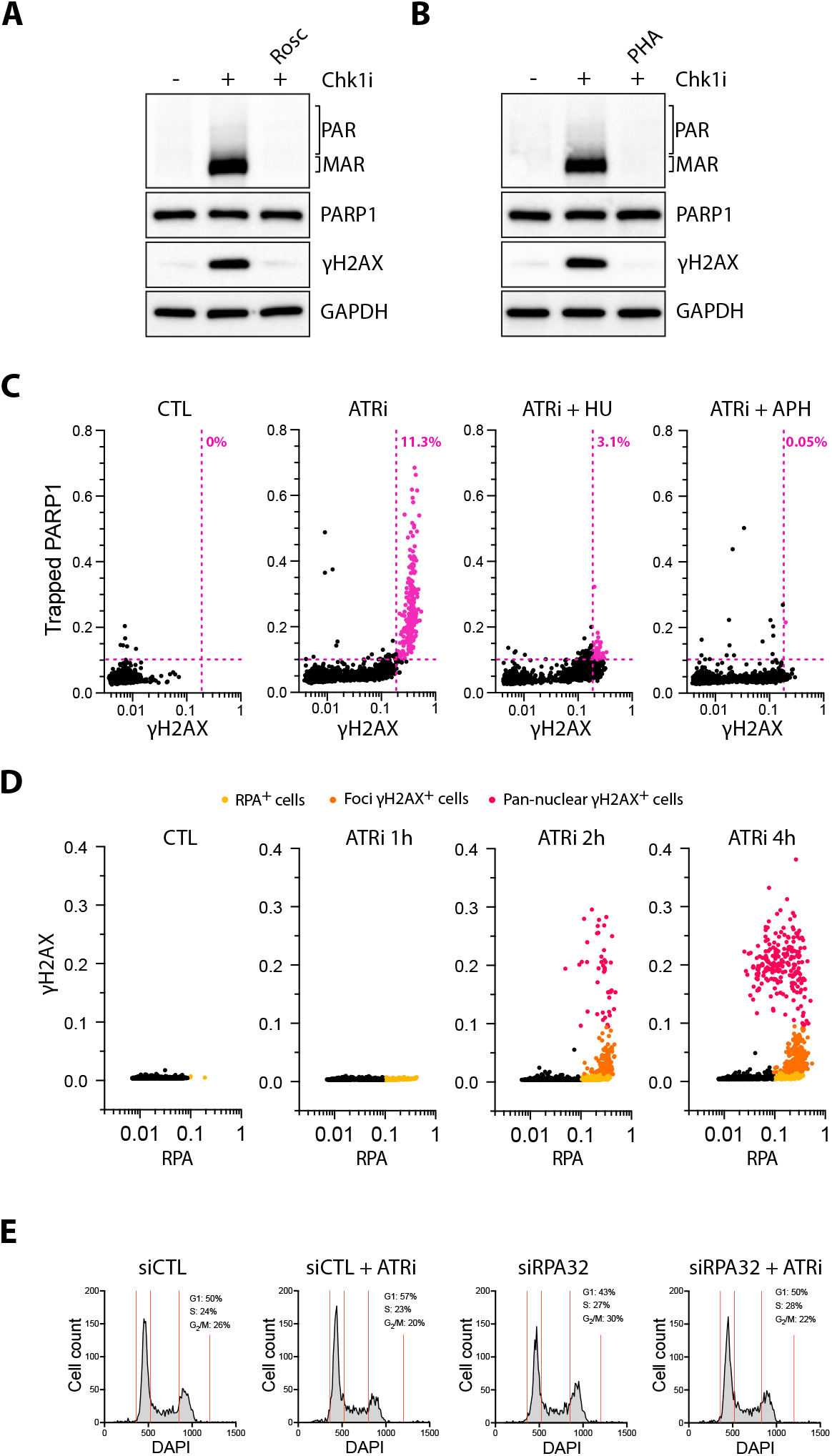
**A-B** The levels of poly/mono ADP-ribose, PARP1, γH2AX, and GAPDH were analyzed by western blot in U2OS treated with Chk1i (1 μM) for 4h in the presence or absence of Roscovitine (12.5 μM) **(A)** or PHA-793887 (3 μM) **(B). C**. Quantification by immunofluorescence of chromatin-bound PARP1 and γH2AX levels in 2,000 U2OS cells treated with ATRi (1 μM) ± hydroxyurea (2 mM) or aphidicolin (0.25 μg/mL) for 4h. Colored dots and percentages indicate cells positive for γH2AX and PARP1. **D**. U2OS cells were treated with ATRi (1 μM) for 1, 2, or 4h. Cells were then analyzed by immunofluorescence against RPA and γH2AX (number of cells, n=2,000). Cells were color-coded as follows: yellow for RPA-positive cells only, orange for RPA-positive cells with low γH2AX intensity, and red for RPA-positive cells with high γH2AX intensity. **E**. Cell cycle analysis by immunofluorescence of U2OS cells transfected with siCTL or siRPA32 for 40h and subsequently treated with ATRi for 4h.

**Supplementary Figure 3:**
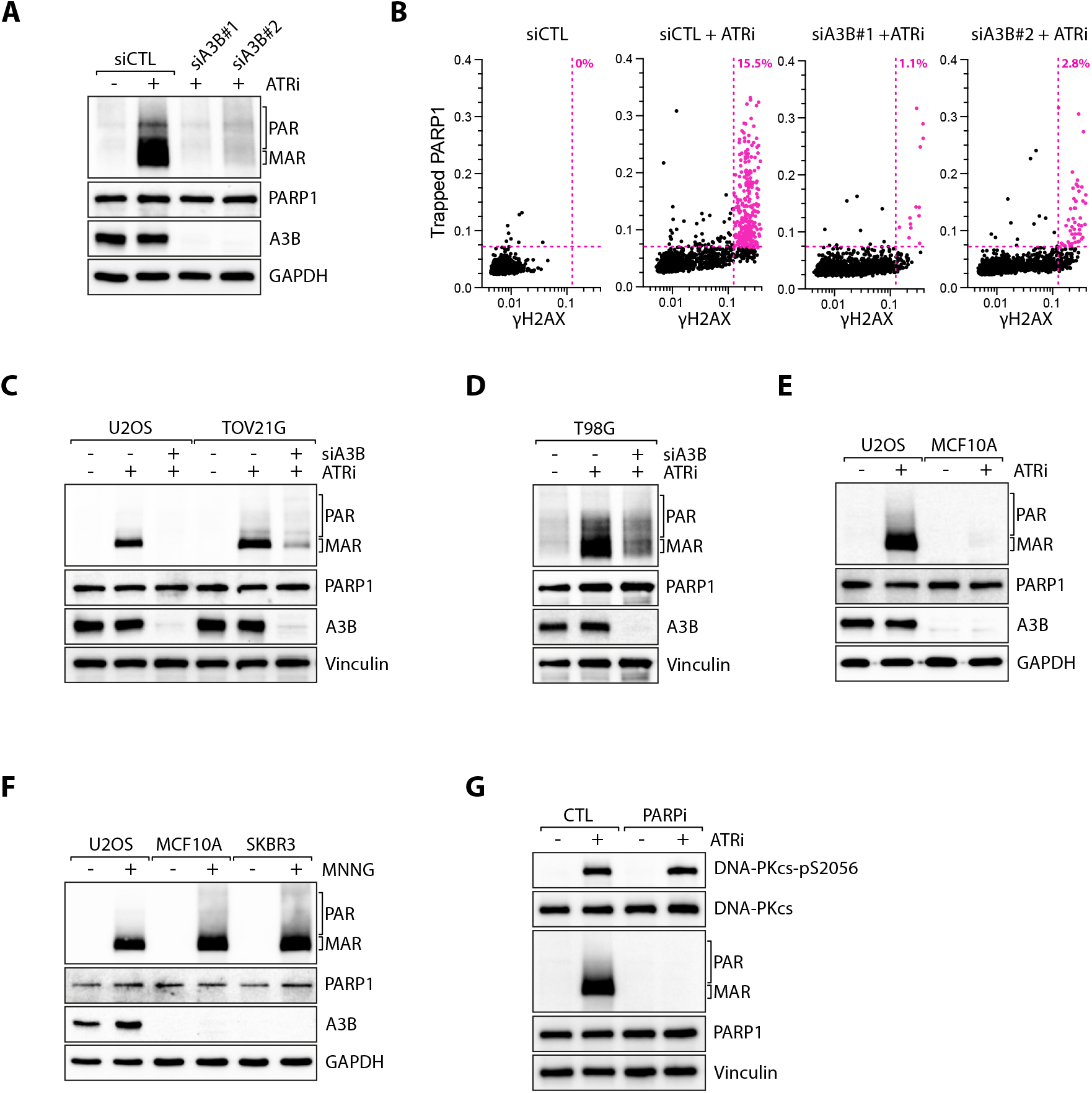
**A-B** U2OS cells were transfected with a siRNA control (CTL) or against A3B for 40h and subsequently treated with ATRi (1 μM) for 4h. Cells were then analyzed by western blot with antibodies against the indicated proteins **(A)** or by immunofluorescence against PARP1 and γH2AX (number of cells, n=2,000) **(B)**. Colored dots and percentages indicate cells positive for γH2AX and PARP1. **C-D**. U2OS, TOV21G, and T98G cells were transfected with a siCTL or siA3B for 40h and subsequently treated with ATRi (1 μM) for 4h. The levels of poly/mono ADP-ribose, PARP1, A3B, and Vinculin were detected by western blot. **E**. The levels of poly/mono ADP-ribose, PARP1, A3B, and GAPDH were analyzed by western blot following ATRi treatment (1 μM) for 4h in U2OS or MCF10A cells. **F**. U2OS, MCF10A, and SKBR3 cells were treated with MNNG (10 μM) for 15 min. The levels of poly/mono ADP-ribose, PARP1, A3B, and GAPDH were detected by western blot. G. U2OS cells were treated with ATRi (1 μM) in the presence or absence of PARPi (20 μM) for 4h, and the levels of indicated proteins were analyzed by western blot.

**Supplementary Figure 4:**
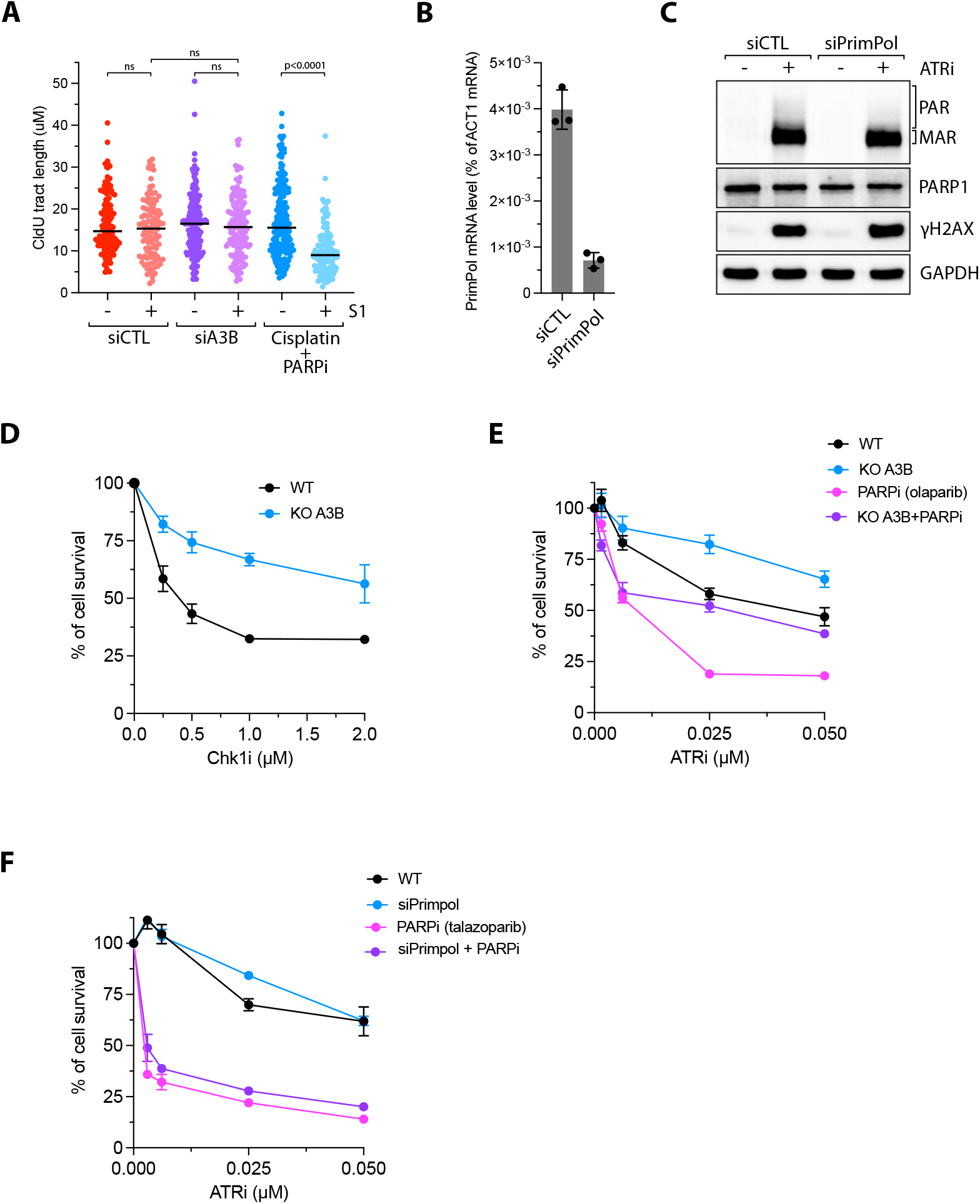
**A**. Quantification of 5-Chloro-2′-deoxyuridine (CIdU) tract lengths in U2OS cells transfected with siCTL or siA3B. When indicated, cells were treated with S1 nuclease before performing the DNA fiber assay. Treatment with cisplatin (150 μM) for 1h and PARPi (100 μM) for 2h serves as a positive control for ssDNA gap-generating conditions. **B**. The levels of PrimPol mRNA were analyzed by RT-qPCR in U2OS cells transfected with siRNA control (CTL) or against PrimPol for 40h. **C**. U2OS cells were transfected with a siCTL or siPrimPol for 40h and subsequently treated with ATRi (1 μM) for 4h. The levels of poly/mono ADP-ribose, PARP1, γH2AX, and GAPDH were detected by western blot. **D**. U2OS WT or A3B KO cells were treated with increasing concentrations of Chk1i for 24 h and then cultured in inhibitor-free media for 48h. Mean values ± SEM. Number of biological replicates, n = 3. **E**. Cell survival of U2OS WT or A3B KO cells treated with increasing concentration of ATRi ± PARPi (1 μM olaparib) for 24 h and then cultured in inhibitor-free media for 48 h. Mean values ± SEM. Number of biological replicates, n = 3. **F**. Cell survival of U2OS transfected with siCTL or siPrimPol for 40h, then treated with increasing concentration of ATRi ± PARPi (1 μM talazoparib) for 24 h, and finally cultured in inhibitor-free media for 48 h. U2OS. Mean values ± SEM. Number of biological replicates, n = 3. All P-values were calculated with a two-tailed Student t-test.

**Supplementary Figure 5:**
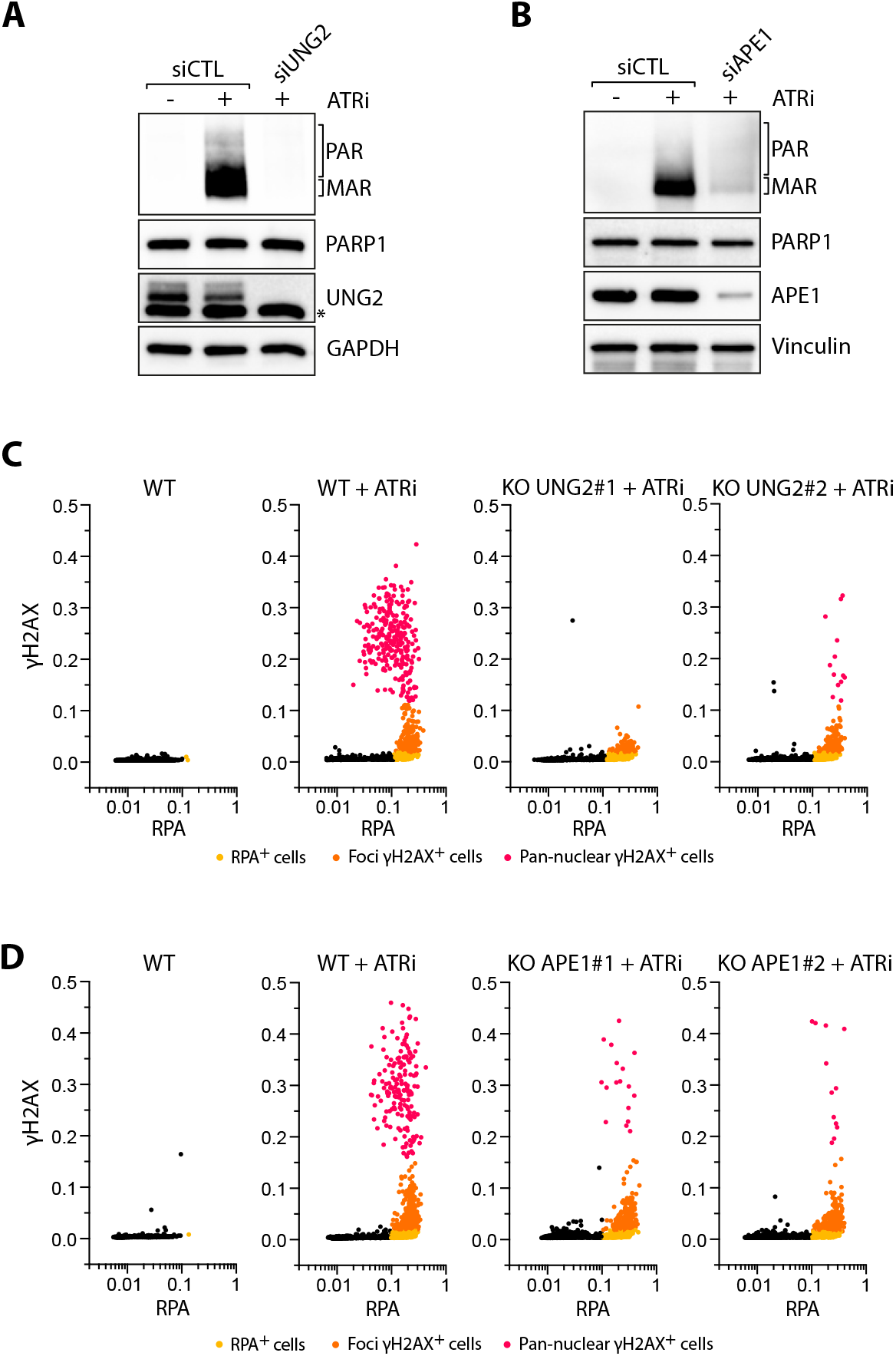
**A**. U2OS cells were transfected with a siRNA control (CTL), against UNG2 **(A)**, or against APE1 **(B)** for 40h and subsequently treated with ATRi (1 μM) for 4h. Cells were then analyzed by western blot with antibodies against poly/mono ADP-ribose, PARP1, UNG, APE1, and GAPDH. **C-D** Quantification by immunofluorescence of γH2AX and chromatin-bound RPA levels of 2,000 U2OS WT, UNG2 KO **(C)** or APE1 KO **(D)** cells treated with ATRi (1 μM) for 4h. Cells were color-coded as follows: yellow for RPA positive cells only, orange for RPA positive cells with low γH2AX intensity, and red for RPA positive cells with high γH2AX intensity.

**Supplementary Figure 6:**
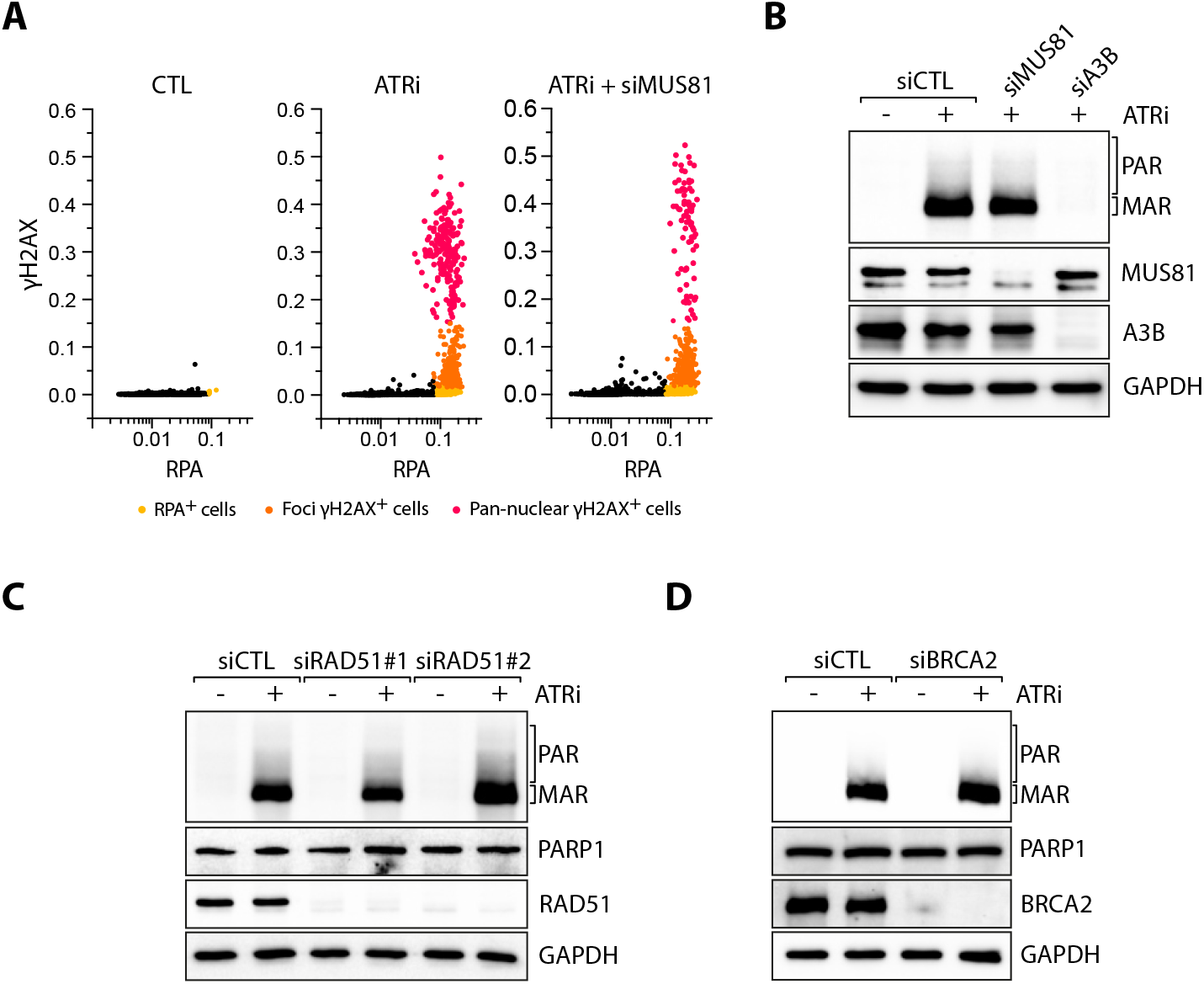
**A**. Quantification by immunofluorescence of γH2AX and chromatin bound RPA levels of 2,000 U2OS cells were transfected with a siRNA control (CTL) or against MUS81 for 40h and subsequently treated with ATRi (1 μM) for 4h. **B**. U2OS cells were transfected with siRNA against MUS81 or A3B for 40h and subsequently treated with ATRi (1 μM) for 4h. The levels of MAR/PAR were analyzed by western blot. **C-D**. The levels of poly/mono ADP-ribose, PARP1, RAD51 or BRCA2, and GAPDH were analyzed by western blot in U2OS cells transfected with siRNA against RAD51 **(C)** or against BRCA2 **(D)** and treated with ATRi (1 μM) for 4h.

**Supplementary Figure 7:**
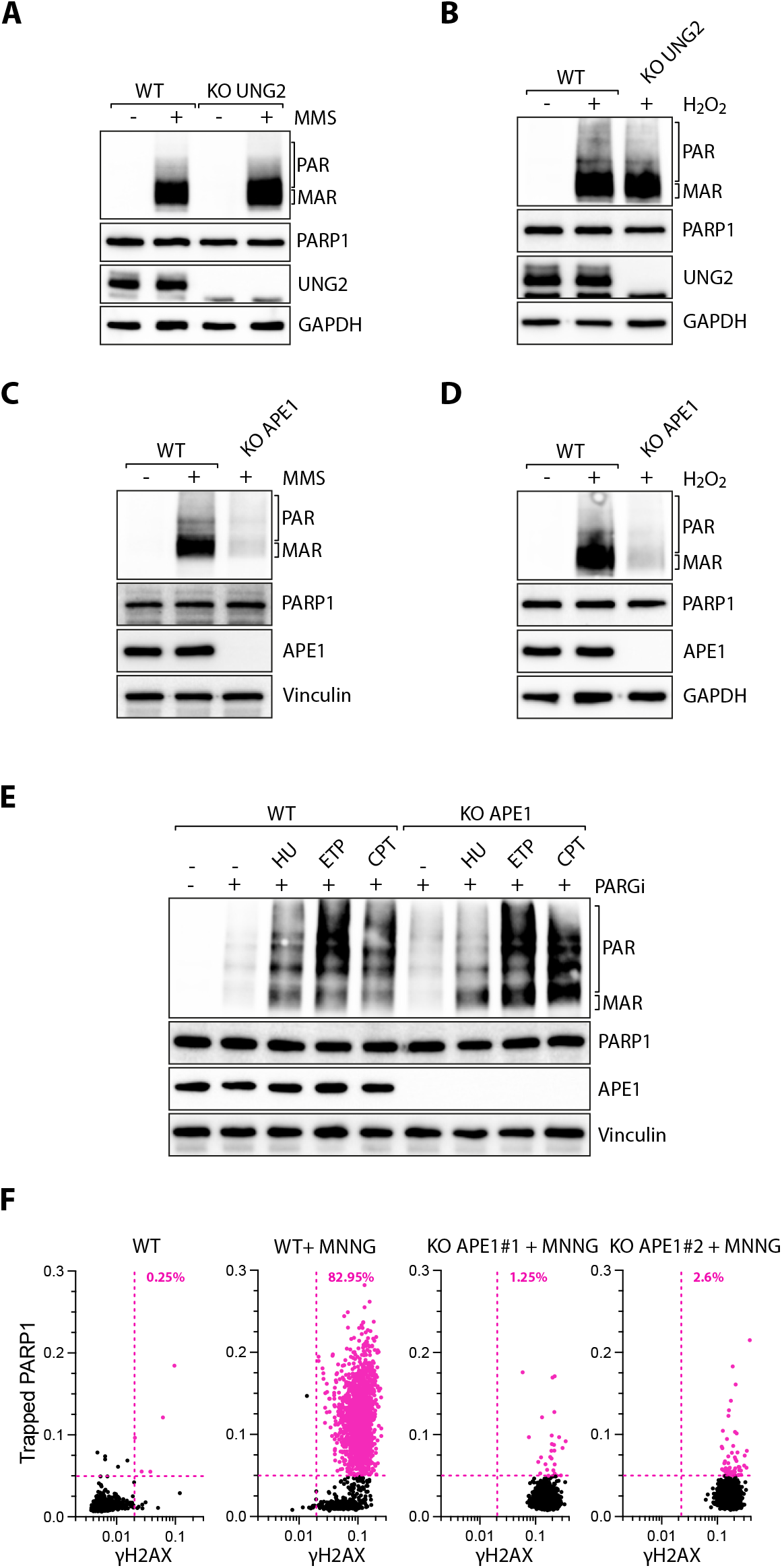
**A-B**. U2OS WT or UNG2 KO cells were treated with MMS (0.01%) for 1h **(A)** or H2O2 for 7 min **(B)** and analyzed by western blot with antibodies against the indicated proteins. **C-D**. The levels of poly/mono ADP-ribose, PARP1, APE1, Vinculin, and GAPDH were analyzed by western blot in U2OS WT or APE1 KO cells treated with MMS (0.01%) for 1h **(C)** or H2O2 for 7 min **(D). E**. U2OS WT of APE1 KO cells were treated with HU (2 mM), ETP (25 μM), or CPT (1 μM) for 4h in the presence or absence of PARGi (2 μM). The levels of poly/mono ADP-ribose, PARP1, APE1, and Vinculin were analyzed by western blot. **F**. U2OS WT or APE1 KO cells were treated with MNNG (10 μM) for 1h and analyzed by immunofluorescence (number of cells, n=2,000) with indicated PARP1 and γH2AX antibodies. Colored dots and percentages indicate cells positive for γH2AX and PARP1.

